# A spatial model of autophosphorylation of Ca^2+^/calmodulin-dependent protein kinase II (CaMKII) predicts that the lifetime of phospho-CaMKII after induction of synaptic plasticity is greatly prolonged by CaM-trapping

**DOI:** 10.1101/2024.02.02.578696

**Authors:** Thomas M. Bartol, Mariam Ordyan, Terrence J. Sejnowski, Padmini Rangamani, Mary B. Kennedy

## Abstract

Long-term potentiation (LTP) is a biochemical process that underlies learning in excitatory glutamatergic synapses in the Central Nervous System (CNS). The critical early driver of LTP is autophosphorylation of the abundant postsynaptic enzyme, Ca^2+^/calmodulin-dependent protein kinase II (CaMKII). Autophosphorylation is initiated by Ca^2+^ flowing through NMDA receptors activated by strong synaptic activity. Its lifetime is ultimately determined by the balance of the rates of autophosphorylation and of dephosphorylation by protein phosphatase 1 (PP1). Here we have modeled the autophosphorylation and dephosphorylation of CaMKII during synaptic activity in a spine synapse using MCell4, an open source computer program for creating particle-based stochastic, and spatially realistic models of cellular microchemistry. The model integrates four earlier detailed models of separate aspects of regulation of spine Ca^2+^ and CaMKII activity, each of which incorporate experimentally measured biochemical parameters and have been validated against experimental data. We validate the composite model by showing that it accurately predicts previous experimental measurements of effects of NMDA receptor activation, including high sensitivity of induction of LTP to phosphatase activity *in vivo,* and persistence of autophosphorylation for a period of minutes after the end of synaptic stimulation. We then use the model to probe aspects of the mechanism of regulation of autophosphorylation of CaMKII that are difficult to measure *in vivo*. We examine the effects of “CaM-trapping,” a process in which the affinity for Ca^2+^/CaM increases several hundred-fold after autophosphorylation. We find that CaM-trapping does not increase the proportion of autophosphorylated subunits in holoenzymes after a complex stimulus, as previously hypothesized. Instead, CaM-trapping may dramatically prolong the lifetime of autophosphorylated CaMKII through steric hindrance of dephosphorylation by protein phosphatase 1. The results provide motivation for experimental measurement of the extent of suppression of dephosphorylation of CaMKII by bound Ca^2+^/CaM. The composite MCell4 model of biochemical effects of complex stimuli in synaptic spines is a powerful new tool for realistic, detailed dissection of mechanisms of synaptic plasticity.

## Introduction

Memories are stored in the brain through creation of new neural networks that are formed by strengthening excitatory glutamatergic synapses between neurons that are activated together during an experience (1–3). Each excitatory pyramidal neuron in the forebrain contains approximately 10,000 excitatory glutamatergic synapses arrayed along several dendrites that reach into the surrounding brain tissue. Most excitatory synapses are comprised of a release site from a presynaptic axon that makes a specialized synaptic contact with a postsynaptic spine, which is a small tubular membrane extension with a bulbous head (Fig. 1A). The spine contains highly integrated biochemical machinery that regulates synaptic strength by controlling the size of the spine head and the number of α-amino-3-hydroxy-5-methyl-4-isoxazolepropionic acid (AMPA)-type glutamate receptors (AMPARs) located at the synaptic site (4–7). Activation of N-methyl-D-aspartate-type glutamate receptors (NMDARs) at synapses in the CNS triggers changes in synaptic strength that underlie memory formation in response to strong synaptic stimuli (8, 9). Precise kinetic control of this machinery determines the amplitude of changes in synaptic strength that are the basis of new neural networks.

**Figure 1.**
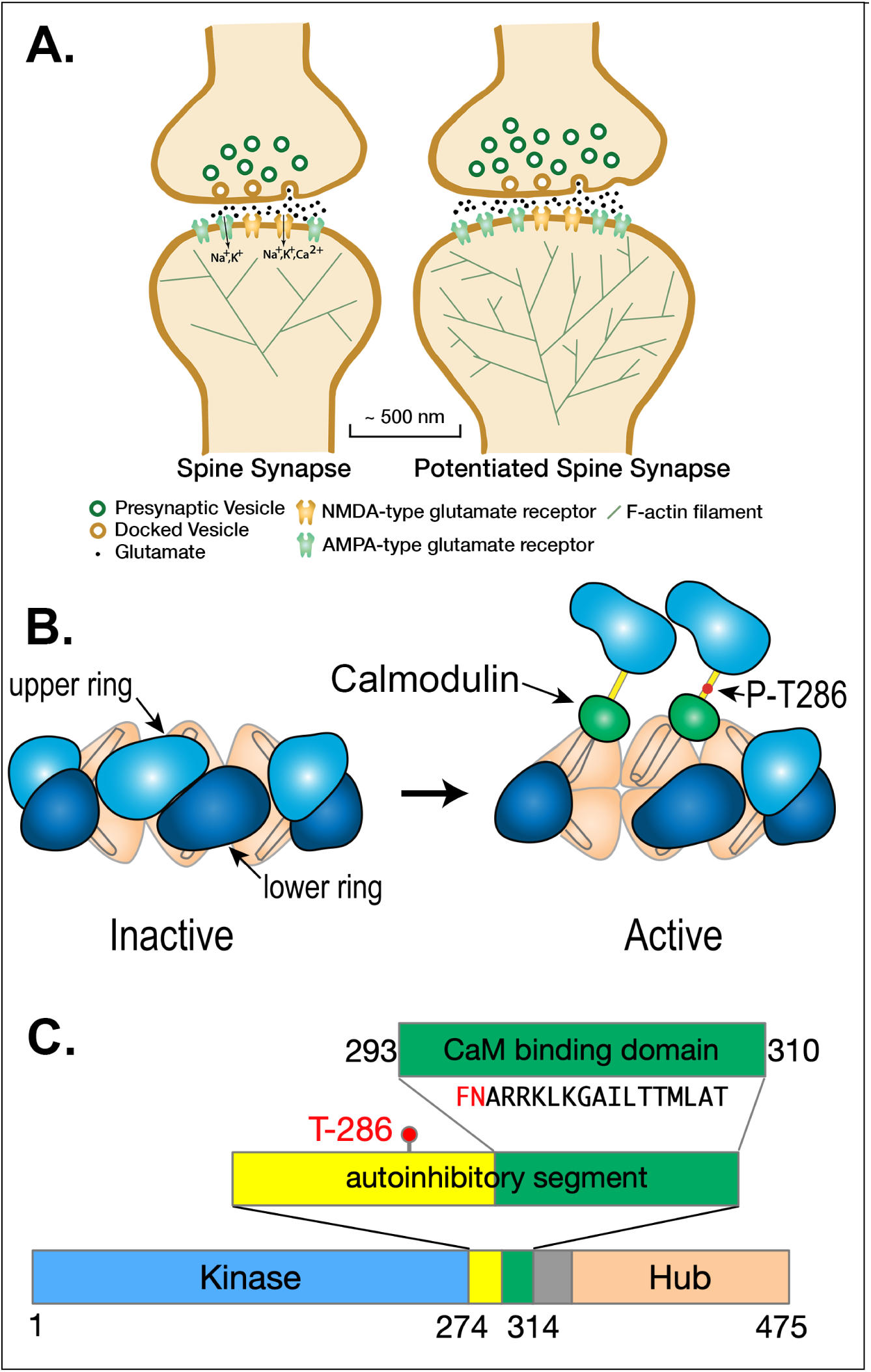
**A**. Schematic diagram of a spine synapse. AMPA-type glutamate receptors (AMPARs, teal) produce a depolarizing Na^+^/K^+^ current upon binding glutamate released from presynaptic vesicles after arrival of an axonal action potential (AP). NMDA-type glutamate receptors (NMDARs, gold) contain a channel that is opened by the concurrence of glutamate release and strong depolarization (e.g. a back-propagating action potential [bAP]). Potentiated synapses (right) contain more AMPARs resulting in a larger excitatory postsynaptic potential (EPSP) upon release of glutamate. The spines of potentiated synapses increase in size and contain a more highly branched actin cytoskeleton. **B**. Ca^2+^/calmodulin-dependent protein kinase II (CaMKII) is a dodecameric holoenzyme in which twelve subunits are bound together by the interactions among the hub domains of each subunit. When Ca^2+^/calmodulin (Ca^2+^/CaM) binds to two adjacent subunits in one of the hexameric rings, a threonine residue (Thr286) within one of the two subunits is autophosphorylated by the other subunit acting as a kinase. Autophosphorylation is believed to occur in one direction around the hexameric ring. Autophosphorylation of Thr 286 causes the subunit to remain active even when the Ca^2+^ concentration falls and calmodulin (CaM) unbinds. [modified from Fig. 7 of Rosenberg et al. (15)]. **C**. Domain diagram of a subunit of CaMKII illustrating the catalytic domain (Kinase), the autoinhibitory segment with the CaM-binding domain, and the Hub domain. When the kinase is inactive, the inhibitory segment lies within the site that binds ATP and protein substrates. Binding of Ca^2+^/CaM to the CaM-binding domain moves the inhibitory segment out of the substrate-binding pocket allowing ATP to bind to the subunit. Thr-286, which can be autophosphorylated, as described in B), is contained within the autoinhibitory segment. Autophosphorylation unmasks a phenylalanine (F) and a glutamine (N) in the CaM-binding domain (shown in red), enabling a large increase in the affinity of Ca^2+^/CaM, known as CaM-trapping.

### Spine synapses and LTP

Spines are, on average, ∼1-2 µm in length. The diameters of spine heads are highly variable, as are their volumes (10, 11, see 12). The median volume of spines in stratum radiatum of hippocampal area CA1 (the source of our model spine) is ∼0.017 µm^3^ or ∼17 x 10^-18^ liters (11). The synaptic membrane contacts between axon release sites and spines are roughly circular and their median diameter is ∼200 nm (11). The presynaptic side of the contact contains sites at which vesicles dock at the membrane and release glutamate probabilistically when the axon fires an action potential. The postsynaptic contact is undergirded by the postsynaptic density (PSD), which immobilizes the synaptic glutamate receptors and anchors a dense network of cytosolic regulatory proteins attached to the membrane (7, 10, 13). The strength of a synapse is highly correlated with the volume of the spine head, as well as the area of the PSD and presynaptic active zone (14). Larger spine heads contain more AMPA-type glutamate receptors and a larger PSD. Thus, they exhibit a larger excitatory postsynaptic potential when glutamate is released from the presynapse (11).

The regulatory biochemical machinery in spines is capable of responding to the frequency of synaptic activation by strengthening (enlarging) or weakening (shrinking) the synapse (6, 16). The biochemical mechanisms that allow the frequency of presynaptic activity to be translated precisely into synaptic strengthening or weakening that faithfully encodes memories are still incompletely understood. The reactions that lead to functional and structural changes in spines are triggered by Ca^2+^ entering the spine through the ion channels of NMDA-type glutamate receptors (NMDARs)(Fig. 1A). Physiological studies show that these functional and structural changes are exquisitely tightly regulated by the frequency and amplitude of Ca^2+^ entry (17, 18).

The principal target of Ca^2+^ entering the spines of cortical and hippocampal excitatory synapses is Ca^2+^/calmodulin-dependent protein kinase II (CaMKII), which, as its name suggests, is activated when it binds Ca^2+^-bound calmodulin (Ca^2+^/CaM) formed when the concentration of Ca^2+^ rises above its baseline value (∼100 nM, 19) (Fig. 1B). Activation of CaMKII has been shown to play a crucial role in long term potentiation (LTP) of excitatory synaptic strength in the hippocampus, and therefore in spatial learning and memory (20, 21). The regulation of CaMKII by Ca^2+^/CaM has also been studied extensively *in vitro* (19, 22–25). Activated CaMKII subunits within the holoenzyme become autophosphorylated at Thr 286 within the first several seconds of Ca^2+^ influx, eventually enabling binding of these subunits to the GluN2B subunits of NMDARs (26, 27). Activation of CaMKII by autophosphorylation also enables phosphorylation of several additional synaptic proteins (28–31). These events set in motion a cascade of reactions that lead to changes in synaptic strength, the magnitude of which depend on the frequency of presynaptic action potentials (APs) and the presence of other regulatory modulators that can influence the sensitivity of the synapse to plastic changes (4, 6, 32).

Modeling of the dynamics of synaptic mechanisms, including activation of CaMKII, has been carried out at different levels of abstraction and mechanistic realism to address different questions. Previous models have often employed simplifications of complex spatial mechanisms, and/or included coarse-grained variables to represent a series of linked mechanisms over several time scales. Many of these have been deterministic compartmental models that rely on differential equations assuming well-mixed conditions in linked small compartments (33–36). In one previous modeling study designed to test the limits of the popular hypothesis that CaMKII constitutes a “bistable” switch that would outlast molecular turnover (37), the authors searched for parameters that would produce such a stimulus-dependent “bistable” switch composed of CaMKII and the protein phosphatase PP1 (e.g. 38). However, the parameter ranges they identified as necessary for this form of irreversible “bistability” are not consistent with data regarding the concentrations of these proteins in synaptic spines. Our study also does not predict the existence of a bistable switch composed of CaMKII and PP1 that would outlast molecular turnover.

Here, we have used modeling to study biochemical responses in spine synapses that are dictated by actual biological structures. Although deterministic methods are useful and appropriate for modeling relatively large compartments that are well-mixed, the biochemical reactions that initiate synaptic plasticity *in vivo* occur within extremely small spines which contain small numbers (ten to a few hundred) of spatially-organized signaling proteins (see 13, 39). Thus, the biochemical molecular mechanisms initiated in spines in the first few minutes of intense synaptic activity involve binding and chemical reactions that occur in a discontinuous, stochastic fashion that is not well represented by differential equations. They also occur in spatially organized compartments that do not satisfy the “well-mixed” assumptions required for the use of deterministic methods to model reaction dynamics. The ability of MCell4 to efficiently represent both the spatial organization of molecular interactions in small spaces and the stochastic nature of state changes (40) makes it an especially useful modeling tool for probing the details of synaptic molecular mechanisms that are driven *in vivo* by the fluctuating, probabilistic Ca^2+^ signals that arise in spines during the first few minutes of synaptic activity. For example, the agent-based, “network free” methods employed in MCell4 with BNGL allow us to avoid simplifying assumptions regarding the interactions of Ca^2+^, CaM, and CaMKII (∼18^12^ states) described below. Methods based on differential equations or population-based stochastic methods (e.g. any method based on the Gillespie Stochastic Simulation Algorithm [SSA], 41, 42) would not be able to capture the combinatorial complexity inherent in these interactions without simplifying assumptions.

We constructed a composite model of activation of CaMKII in a spine by combining four previously published models. We first show that the composite model produces the observed behavior of NMDARs and resulting Ca^2+^ influx into a spine. We validate it as a model of synaptic activation of CaMKII by showing that it predicts *a priori* aspects of phosphorylation and dephosphorylation of CaMKII *in vivo* that have been experimentally measured, specifically high sensitivity to protein phosphatase activity (e.g. 43) and persistence of autophosphorylation for a few minutes after the end of the synaptic stimulus (44). We then use the model to test hypotheses about the role of CaM-trapping (described below) in regulation of synaptic plasticity that are not easily testable experimentally.

### Medical Relevance

Modeling of the precise kinetics of synaptic regulatory mechanisms will have medical relevance. Several intractable mental and neurological illnesses, including Alzheimer’s disease (45), involve disruptions in mechanisms of synaptic regulation (46–48) that can produce relatively subtle, but devastating, changes in the timing of synaptic reactions. Given the central role of CaMKII activation for synaptic plasticity, it is important to understand the detailed biochemical dynamics of the Ca^2+^-CaM-CaMKII signaling axis within spines to inform the design of new therapeutics to treat these illnesses.

### Structure of CaMKII

CaMKII is a dodecameric holoenzyme comprised of twelve individual catalytic subunits that are organized via their “hub” domains into two hexameric rings stacked together (Fig. 1B, C; 15, 49, 50). The kinase activity of each subunit is activated independently by binding of Ca^2+^/CaM to its CaM-binding domain, which moves the inhibitory domain away from the catalytic pocket and thus activates kinase activity (51). In addition, the subunits can undergo autophosphorylation in which an active subunit phosphorylates a specific threonine residue (Thr 286) within its neighboring subunit if that subunit also has Ca^2+^/CaM bound to it (22, 52–55). Phosphorylation of Thr 286 blocks inhibition by the regulatory domain (51); therefore, each autophosphorylated subunit stays active as long as Thr 286 remains phosphorylated, and can, in turn, phosphorylate and activate its neighboring subunit when the neighbor binds Ca^2+^/CaM. Thr 286 can be dephosphorylated by either of two broad-specificity protein phosphatases, phosphatase 1 (PP1) or -2a (PP2a); however, PP1 activity predominates in spines (56–61).

In an important study, Meyer and Schulman (23) found that the initial binding of Ca^2+^/CaM to a CaMKII subunit occurs with moderate affinity (K_D_ = ∼50 nM). However, the movement of the inhibitory segment caused by autophosphorylation substantially increases the affinity for Ca^2+^/CaM, a process they referred to as CaM-trapping. Putkey and Waxham (62) later found that the off-rate of CaM is increased from 6.6 s^-1^ to 9x10^-5^ s^-1^, producing an ∼70,000-fold increase in affinity (K_D_ = ∼1.8 pM).

Tse et al. (63) showed that autophosphorylation exposes residues Phe293 and Asn294 in the CaM-binding domain, and these residues mediate the high affinity CaM-trapping. After autophosphorylation, residues on both the N- and C-termini of CaM (Fig. 2A) bind tightly around the FNARRK sequence in the CaM-binding domain (Fig. 1C, Fig. 2).

**Figure 2.**
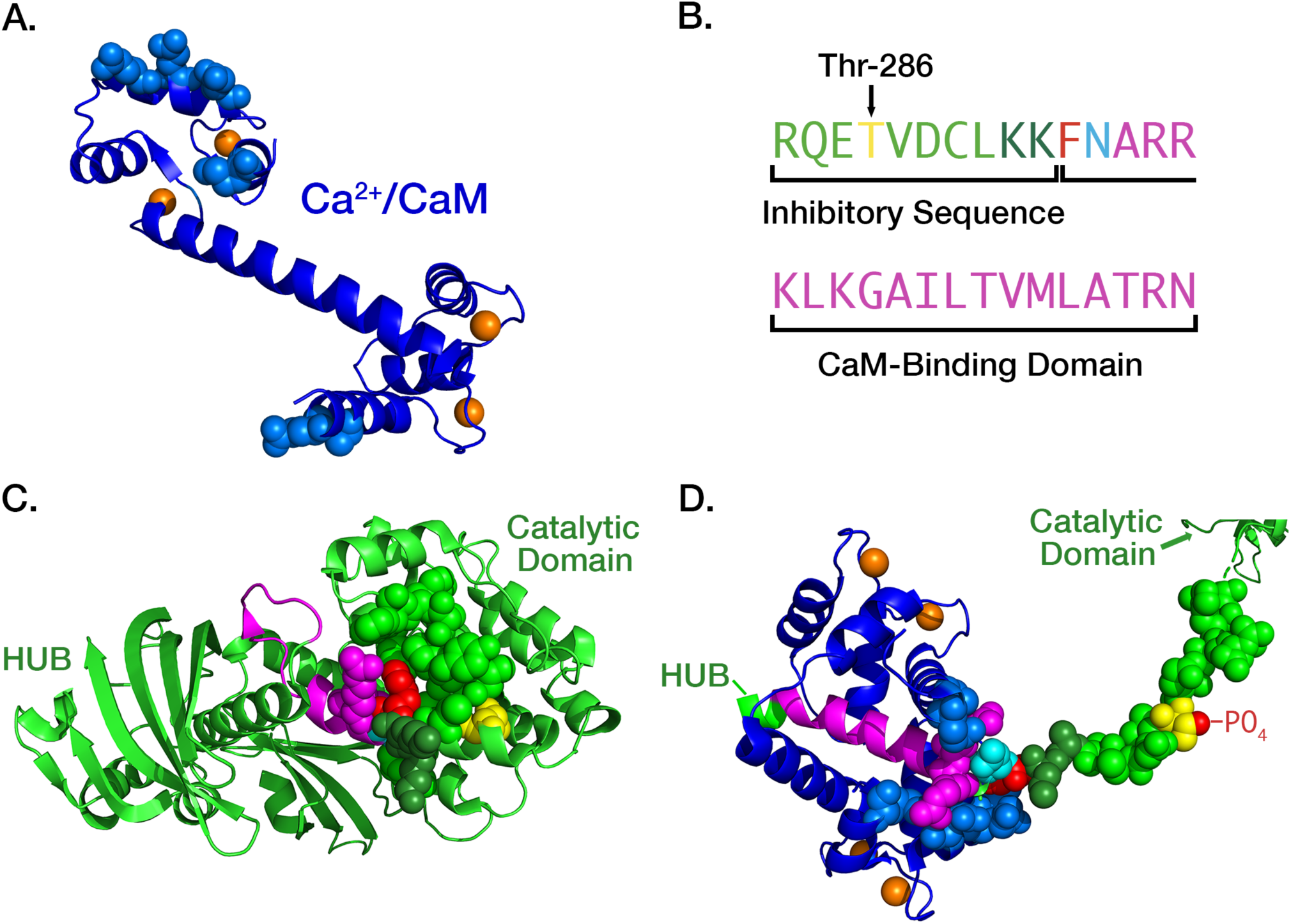
A. Ribbon diagram of the atomic structure of CaM with four bound Ca^2+^ ions (orange), two each at its N- and C-termini. The atoms of residues that are bound tightly to the inhibitory segment of CaMKII during trapping are represented as spheres. Rendered in PyMol from pdb #1CLL; B. Color-coding of residues comprising the inhibitory and CaM-binding domains of CaMKII subunits: Thr 286 (yellow), two K residues shielding the F and N residues (dark green), F (red), N (cyan), initial CaM-binding domain (magenta). C. Ribbon diagram of the atomic structure of an autoinhibited subunit of CaMKII (light green) with the atoms of residues that mask the CaM-trapping amino acids phenylalanine (F, red) and asparagine (N, cyan) shown as light and dark green spheres. Rendered in PyMol from pdb #3SOA; D. Ribbon diagram of “trapped” CaM bound to the extended CaM-binding domain. After autophosphorylation of Thr-286 the inhibitory segment assumes a fully extended conformation. The critical F and N residues are exposed and residues on both the N- and C-termini of CaM (blue) bind tightly around the FNARRK portion of the CaM binding domain. Rendered in PyMol from pdb #2WEL.

Various investigators have postulated that CaM-trapping leads to an enhanced non-linear increase in the rate of autophosphorylation of CaMKII during high frequency synaptic stimulation (9, 64, 65). However, the tiny spatial compartment of the spine, and the low copy numbers of molecules therein make it impossible to measure enzyme activity experimentally within spine compartments without significantly perturbing the kinetics of protein-protein interactions. Computational models can act as a kind of kinetic “microscope” to explore how detailed dynamics occur within small cellular compartments. They also permit tests of the importance of individual components of a model, by revealing the consequences of the experimental measurements built into it, and of variations of the model’s parameters. Here, we have considered how CaM-trapping will affect the activation and autophosphorylation of CaMKII holoenzymes in a median-sized spine responding to a presynaptic stimulus consisting of two 1 sec bursts of 50 Hz presynaptic stimulation, spaced 2 sec apart. Under these conditions, we show that CaM-trapping does not influence the proportion of autophosphorylation of CaMKII during the intermediate-strength two-epoch stimulus because it does not increase the number of non-phosphorylated subunits with bound CaM; therefore, it doesn’t increase the probability of new autophosphorylation during the second stimulus epoch. We show that this is the case whether the holoenzymes are distributed uniformly through the spine or are concentrated within the PSD.

Instead, we predict that CaM-trapping will prolong the lifetime of autophosphorylated CaMKII after a stimulus by as much as an order of magnitude, if, as recently proposed, CaM binding to CaMKII sterically hinders binding of phosphatases to autophosphorylated CaMKII, slowing the rate of dephosphorylation (see 66). We suggest that *in vitro* experiments with purified proteins to measure the rate of dephosphorylation of autophosphorylated CaMKII by PP1 in the presence and absence of Ca^2+^/ CaM will test whether a key effect of CaM-trapping is to slow dephosphorylation of Thr286.

## Methods

### Model development

We modeled activation of CaMKII after a complex presynaptic stimulus delivered to a realistic spine synapse. To do this, we integrated four previous well-tested models to enable us to simulate the activation of dodecameric holoenzymes of CaMKII in a median-sized spine. The models were integrated using the latest version of MCell (MCell4) (40) which permits spatial modeling of reactions specified in the efficient BNGL language (67). The previous models include a model of presynaptic release of glutamate from a CNS synapse (68, 69), a model of Ca^2+^ influx through NMDAR’s and L-type Ca^2+^ channels followed by recovery in reconstructed hippocampal postsynaptic spines and dendrites (12), a model of activation of CaMKII subunits by Ca^2+^/CaM (19), and a model of autophosphorylation of the dodecameric CaMKII holoenzyme (70). Each of these models and their parameters were independently validated against experimental data before they were published. As stated in the introduction, the composite model was not constructed to test particular theories, rather it was built to study as closely as possible the behavior that is dictated by actual biological structures. Accordingly, there are essentially no “free variables” that would represent simplifying assumptions. The numbers of molecules and their spatial locations are based upon multiple examples and forms of experimental data, described here and in the original publications. We have not attempted to model here the full range of influences that regulate CaMKII activity in the spine synapse because such an analysis will require several manuscripts of this size; rather we present an initial model that provides first-order quantitative information about how the CaMKII holoenzyme in a spine would respond during the first 10-20 seconds of activation of NMDARs by a complex stimulus. In future studies, we will be able to add additional elements, including other CaM-binding proteins, in a step-by-step fashion to determine how they influence the dynamics of activation of CaMKII (e.g. 70, 71); its binding to downstream proteins, including GluN2B (26, 27); and its action on enzymatic targets (e.g. 72, 73). Thus, the model will be a powerful tool to study precise biochemical mechanisms of synaptic plasticity by simulating the exact kinetics of the intricate reactions triggered in the first minutes of Ca^2+^ influx through NMDARs.

### MCell 4 Platform

The models are built with the updated agent-based modeling platform MCell4 (40) which integrates the rule-based, network-free simulation framework of BioNetGen (74) with the spatial simulation capabilities of MCell (75, 76). This version builds on the earlier MCell-R, which employed the reaction methods of NFsim originally developed as a non-spatial simulation platform to read and compute simulations written in the model specification language BNGL (67). NFsim’s library of functions carries out the graph operations required for efficient network-free simulation in a non-spatial context. MCell-R extended these operations to encompass the spatial context. Thus, it avoids the need to pre-compute a full reaction network and store each possible state in memory. Instead, it executes each rule as needed. The number of rules specified in BNGL is usually much lower than the number of possible reactions in the full network; thus, memory savings are substantial (76). MCell4 improves upon the performance and generality of MCell-R by replacing the NFsim library with a newly developed library of functions, called libBNG, which implements full integration of network-free BNGL directly in MCell4. The number of possible reactions of the CaMKII holoenzyme described below would be computationally intractable in other modeling platforms. A Python API was added to MCell4 enabling the model presented here by permitting customizations to the configurations of molecules and reactions that are not easily encoded in CellBlender, the graphical user interface for MCell (77).

The workflow for modeling and computation of the full sequence of presynaptic stimulus, glutamate release, NMDAR activation, back-propagating action potential (bAP), Ca^2+^ flux, and autophosphorylation of CaMKII is diagrammed in Fig. 3.

**Figure 3.**
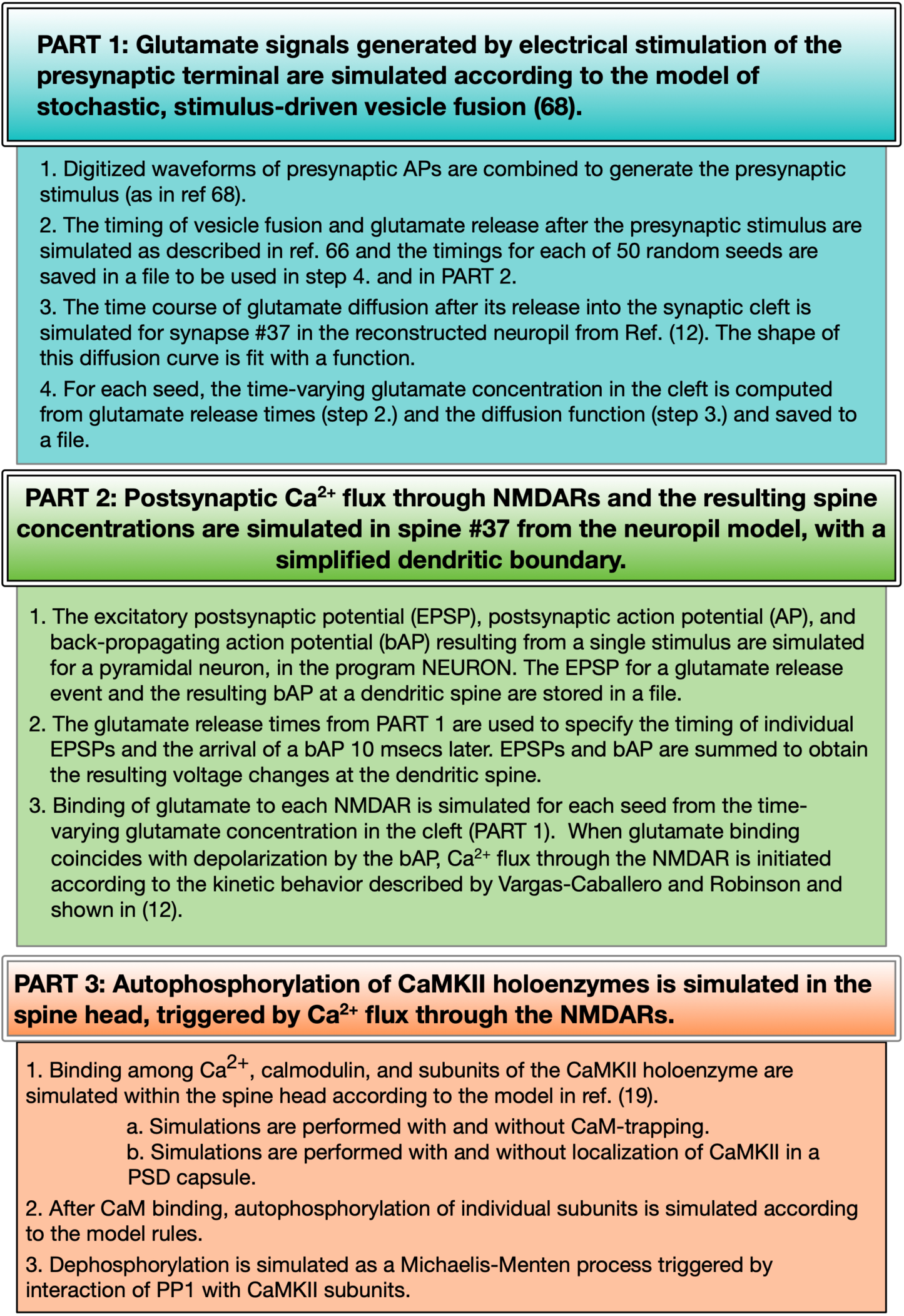
<u>Workflow for simulations with the composite model</u>. In Part 1, the glutamate signals are computed separately for 50 seeds and saved to files. In Part 2, the postsynaptic EPSPs and bAPS at the spine are computed separately in NEURON and saved to files. For each individual simulation, steps 2 and 3 of Part 2, and all of Part 3 are computed in MCell and the changes in state of selected species are saved to output files for analysis and plotting.

### Presynaptic Stimulus

We used a stimulus consisting of two epochs of presynaptic action potentials.

Each epoch consisted of five bursts of action potentials (APs) delivered sequentially at 5 Hz (e.g. 5 bursts per sec). Each burst contained 5 action potentials delivered at 50 Hz (e.g. 5 APs per 100 msec). The two epochs were separated by 2 sec (Fig. S1 A, B). The AP waveforms were generated as described in Ref (68).

### Glutamate Release

Glutamate releases from the presynaptic terminal during the stimulus were simulated based upon our previous MCell model of release from presynaptic vesicles (68, 69). The model of release includes 37 measured parameters, adjusted to 34° C. The model was verified against published experimental studies including: 1) the time course of change of release probability after a single AP (78); 2) magnitude and time course of paired-pulse facilitation (79); and 3) time course of recovery of the release probability after vesicle depletion (69).

A set of 50 realistic stochastic timings of release of glutamate from presynaptic vesicles driven by the two-epoch stimulus were computed in MCell initiated by 50 random seeds. The timings of glutamate release computed in this model took into account stochastic short term plasticity of vesicle fusion, including facilitation by residual Ca^2+^ and depression due to depletion of docked vesicles and resulting decreases in release probability. We assumed a median vesicle size containing 1500 molecules of glutamate (See Fig. 6 in Ref. 80). Releases did not occur with each of the 50 simulated presynaptic APs because for this median-sized spine we simulated the initial release probability as ∼0.2. The active zone initially contained 7 docked vesicles. Release was facilitated after the first few stimuli (due to residual presynaptic Ca^2+^ in the active zone) and then depressed due to the reduction in the number of docked vesicles. Simulations initiated by each seed resulted in different stochastic trains of release times. Empty vesicle docking sites were reoccupied with a time constant of 5 seconds (i.e. 0.2 per second rate of redocking per site).

The precise timing of the Glu release after a presynaptic AP was stochastic and sometimes occurred 10’s of ms after the AP, especially after a long train of stimuli when the [Ca^2+^] in the presynaptic terminal was high (Fig. S1 C, D). If the vesicle docking sites were highly depleted and [Ca^2+^] was elevated in the terminal, a vesicle sometimes appeared to release “spontaneously” at the moment it docked. The sets of timings of glutamate release were indexed by their seed numbers and stored in files in a directory named “glu_release_times_stp_5at50Hz_5at2Hz_2x” and used in the python script, “glu_dependent_rates.py” embedded in the model.

### Concentration of glutamate in the cleft

The full spatial model of hippocampal neuropil described in (12) was used to measure the time courses of glutamate concentration in the synaptic cleft at the location of the cluster of 15 NMDA receptors (see below). Each vesicle fusion released 1500 molecules of Glu and the vesicle release sites were assumed to be located over the NMDAR cluster. NMDAR clusters are usually located immediately adjacent to AMPAR clusters (81), which have been found to be coupled to presynaptic release sites to form transynaptic “nanocolumns” (7). Placement of release sites directly over the NMDAR cluster will not have a large effect on glutamate binding because the on-rate of glutamate binding to NMDARs is much slower than glutamate’s rate of diffusion (82). However, placing the release sites directly over the NMDAR cluster means that these simulations did not reflect the small variability in glutamate concentration at NMDARs that would result from stochastic locations of nanocolumns and release sites. To measure the glutamate concentration, a measurement box was positioned in the cleft covering the extracellular domains of the receptors. The box was transparent to the diffusion of glutamate. Its depth was equal to the height of the cleft and the sides of the box enclosed the NMDAR cluster. The time course of concentration of glutamate within the box was recorded for a series of single releases and then averaged. The averaged transient of glutamate in the box was brief with a decay time constant, *τ*, of about 1.2 µs, and had a peak concentration of ∼70 mM. The time-varying glutamate concentration in the cleft in response to the complex stimulus was computed by summing together the individual glutamate transients occurring at each release time. The resulting concentration time course was used to create a pseudo-first order approximation of binding of glutamate to NMDARs, as follows. If k_plus_ is the second order rate constant for glutamate binding to NMDARs, the rate of binding of glutamate is given by [glu] x [NMDAR] x k_plus_. Binding of glutamate to each NMDAR was specified as a first-order transition from an unbound NMDAR to an NMDAR with bound glutamate. The measured time course of the glutamate concentration was used to compute pseudo-first order rate constants, k_plus_^-^effective = [glu] x k_plus_, for each release event and for each seed, as specified in the python script, “glu_dependent_rates.py.” The k_plus_-effective at each time was then used to determine the probability of transition of each NMDAR to the bound state for each seed during the simulation.

### NMDAR Response and Ca^2+^ Handling in the Spine

The response of NMDARs was modeled with the kinetics used in our earlier MCell model of Ca^2+^ influx into the spine through NMDARs (12, 83). The spine model was originally constructed with MCell3 within the geometry of a hippocampal dendrite reconstructed from serial sections of a 6 µm x 6 µm x 5 µm cube of neuropil imaged by electron microscopy (12). The model includes all of the major sources and sinks of Ca^2+^ ion known to be present in the membranes and cytosol of spines and dendrites. It incorporates experimentally measured kinetics of Ca^2+^ influx through Ca^2+^ channels and NMDARs, including desensitization and the flickering removal of the Mg^2+^ block by a back-propagating action potential. It also includes experimentally measured parameters of Ca^2+^ buffering, and Ca^2+^ removal from the spine via Ca^2+^ exchangers, pumps, and diffusion. Most of the 85 parameters in the model were taken from experimental literature; some were derived from the thermodynamic requirement for microscopic reversibility. All parameters were adjusted to a temperature of 34-37° C. Predicted Ca^2+^ fluxes induced by excitatory postsynaptic potentials (EPSPs) and back-propagating action potentials (bAP’s) simulated in the model were validated against an experimental study by Sabatini et al. (84). For the model presented here, one cluster of 15 NMDARs (Fig. 4) was added near the center of the PSD on the reconstructed spine membrane, reflecting recent data about the number and arrangement of NMDARs in hippocampal synapses (e.g. 81, 85). The positions of other molecules in the spine are documented in detail in the original publication (12).

**Figure 4.**
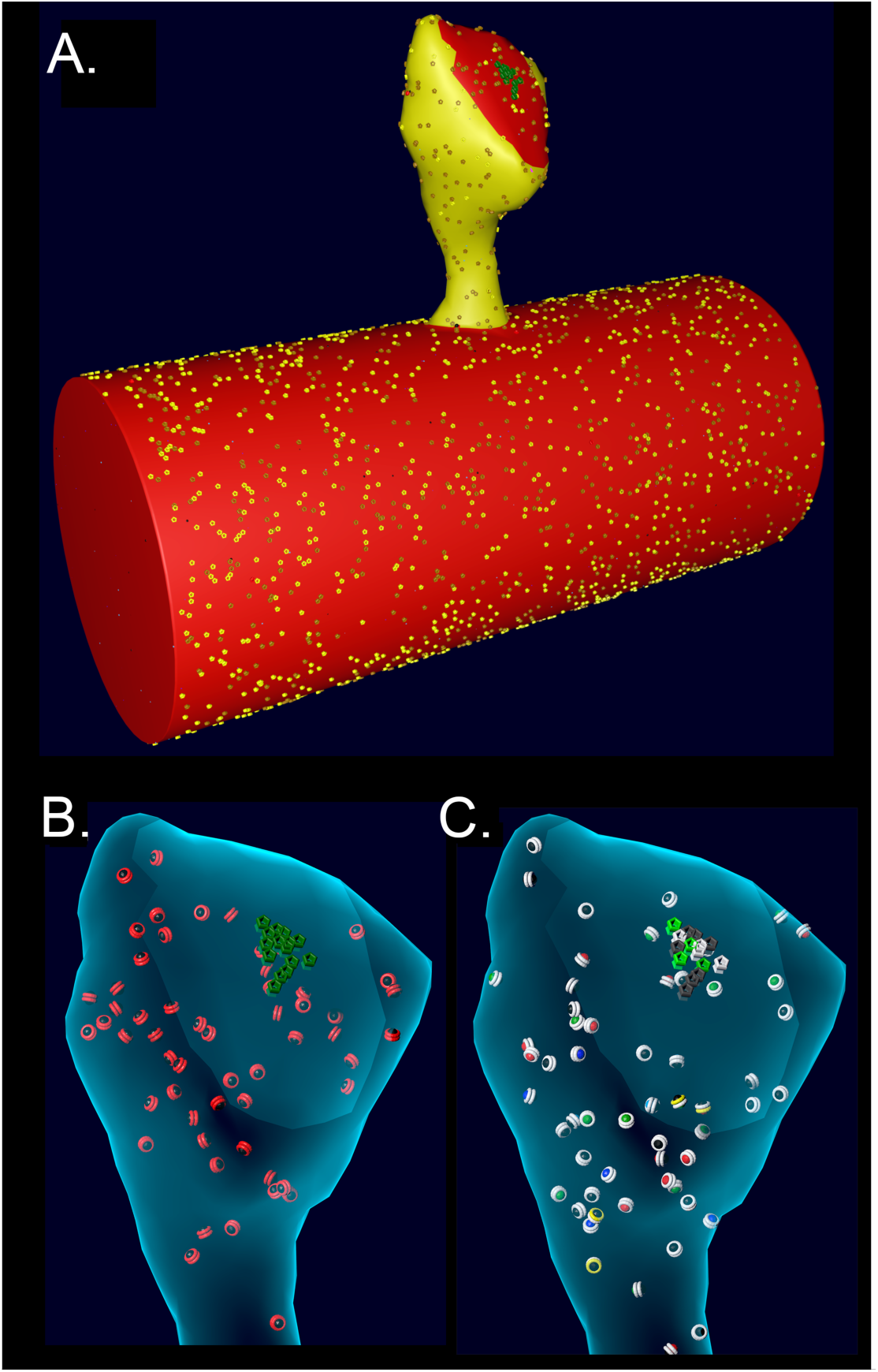
**A**. Spine # 37 from (12) attached to a cylinder with visible glyphs representing surface proteins. The PSD area on the spine and the attached cylinder are shown in red. **B**. A translucent view of spine #37 at instantiation showing surface NMDARs (green) and internal inactive CaMKII holoenzymes (red glyphs). **C**. Spine #37 at t=159 msecs after the start of a stimulus. Open NMDARs on the surface are represented as white glyphs The outer rings of the CaMKII glyphs have changed color to white indicating where CaM with 4 bound Ca^2+^ ions has bound to the subunits (see Results). The color of the internal spheres in the glyphs indicates the number of autophosphorylated subunits in each ring.

### Spine Geometry

To create an accurate model of the response of CaMKII to Ca^2+^ influx inside a spine, while also minimizing computational time, we imported the geometry of a median-sized spine (#37) from the reconstruction in Bartol et al. (12) and attached it to a cylinder 0.7 µm in diameter (84) and 1.5 µm long, created in Blender (Fig. 4). The wall of the spine has the shape of the inner membrane leaflet, and is reflective to all the volume molecules. The mesh representing the attachment of the spine to the cylinder was adjusted to make it free of leaks (i.e. “water-tight”). The volume of the cytosol of the spine is 0.016 µm^3^ (0.016 fl) and the PSD diameter is ∼200 nm. The cylinder and spine are instantiated with the appropriate numbers of Ca^2+^ pumps and channels (NCX, PMCA, VDCC) in the membrane, and with cytosolic molecules Ca^2+^, calbindin, and immobile buffers, as described for the spine and dendritic shaft in (12). We added CaMKII holoenzymes to the cytosol of the spine, encoded as described below, at a concentration of 6.67 µM (60 holoenzymes); thus, individual subunits were present at a concentration of 80 µM (720 subunits, 49, 86). PP1 was added to the spine cytosol at a “baseline” concentration of 1.25 µM (12 molecules), reflecting experimental measurements of the approximate concentration of PPI in neuronal cytosol and its enrichment in spines (87). The k_cat_ for PP1 was set to 11.5/sec (58) and the K_M_ for binding to CaMKII was 11 µM (88). Dephosphorylation was modeled as a Michaelis/Menten reaction (see Parameter Table in Supplementary Methods) triggered stochastically by binding of PP1 to a CaMKII subunit. CaM was added to the cytosol of both the spine and the cylinder at a concentration of 30 µM (89, 90), which is 290 CaM molecules in the spine and ∼10,300 in the cylinder. To test the effect of varying the concentration of PP1 on autophosphorylation, the concentration of PP1 in the spine was varied from 0.65 µM to 5 µM. Details of the calculations of numbers of CaMKII, CaM, and PP1 are in Supplemental Methods.

### Boundary Conditions

In the previous model of Ca^2+^ flux (12), the spines were attached to a reconstructed dendrite. Diffusing molecules, including Ca^2+^ and calbindin, moved freely through the neck between the spine and the dendritic cytoplasm. The VDCCs, PMCAs, and NCXs in the dendritic membrane contributed to the decay time of the spine Ca^2+^ transient. In that model, we showed that in the first 100 msecs after a stimulus, approximately half of the Ca^2+^ entering spine #37 after an EPSP followed by a bAP exited by diffusion through the neck, either free or bound to calbindin, consistent with (84). Thus, in the model presented here, Ca^2+^, calmodulin, and calbindin were allowed to move freely between the spine and the cylinder, but CaMKII holoenzymes and PPI were confined to the spine to facilitate simulation and counting of transformations of CaMKII and its dephosphorylation by PP1 in the spine. To implement this, we constructed a “spine shell” consisting of a mesh encasing the spine at a uniform distance of 20 nm from the spine membrane. The shell cuts through the interface at the junction between the spine and the cylinder so that it forms a closed barrier at the base of the spine neck. CaMKII holoenzymes and PP1 are confined to the spine by making this shell reflective to CaMKII and PP1, but transparent to diffusing Ca^2+^, calmodulin, and calbindin. The dynamics of Ca^2+^, CaM, and CaMKII are quantified within the spine shell.

The length of the cylinder was enlarged until the results of simulations of activation of CaMKII within the spine converged when we enlarged it further. Convergence was achieved between cylinder lengths of 1.5 and 1.8 µm. Therefore, the length of the cylinder was set at 1.5 µm, ensuring that the cylinder provides a boundary equivalent to the dendritic shaft.

### Calmodulin

The model of initial binding interactions among Ca^2+^, calmodulin, and individual subunits of CaMKII was based upon our previous non-spatial model of the dynamics of activation of monomeric CaMKII subunits by Ca^2+^/CaM as the concentration of Ca^2+^ rises (19). This model was created in Mathematica and includes states of Ca^2+^/CaM with one, two, three, or four bound Ca^2+^ ions. The 55 parameters for Ca^2+^ binding to free CaM and for Ca^2+^ binding to CaM bound to CaMKII were measured experimentally at 30° C by various investigators, including the Kennedy lab (91), or were calculated from the measured parameters using the thermodynamic requirement for microscopic reversibility, involving “detailed balance” of ΔG within reaction loops (see 19).

In the present model, the CaM molecule was encoded in BNGL as follows: CaM(C∼0∼1∼2, N∼0∼1∼2, camkii). In BNGL syntax, the name of the molecule is followed in parentheses by a set of “components” demarcated by ∼, which represent binding sites or modification sites. As shown in Fig. 2A, CaM has four binding sites for Ca^2+^, two at the C-terminus and two at the N-terminus; thus, the three states for each terminus (∼0,∼1, or ∼2 of components C or N) represent zero bound Ca^2+^, one bound Ca^2+^, or two bound Ca^2+^, respectively. The camkii component is the binding site for a CaMKII subunit. The model in Pepke et al. (19) did not include CaM trapping following autophosphorylation; CaM-trapping was introduced into the model of the holoenzyme as described below.

### CaMKII Holoenzyme

We updated the model of CaMKII subunits within a dodecameric holoenzyme from Ordyan et al. (70). This non-spatial model employed BioNetGen language (BNGL) and the simulation algorithm NFsim, to build the structure of the holoenzyme, simulate the kinetics of interaction of its subunits with Ca^2+^/CaM, and simulate the subsequent autophosphorylation and dephosphorylation by phosphatase of individual subunits. Parameters and reactions of subunits in this model were based upon those in Pepke et al. (2010), which have been repeatedly vetted for accuracy. This model did not include CaM-trapping or spatial dynamics and was originally used to examine the potential influence of neurogranin on integration and summation of autophosphorylation following increases in Ca^2+^ concentration.

In the present, updated model, the individual subunits are specified in BNGL as follows: <CaMKII(l, r, c, T286∼0∼P, pp1∼0∼1, T306∼0∼P, cam∼la∼ha)> (Fig. 5A). Components l, r, and c are sites of interaction within the holoenzyme; l is a binding site at the left side of the subunit, r at the right side of the subunit, and c (center) a site that interacts with a subunit in the opposite six-membered ring (Fig. 5B). The unphosphorylated subunits were added (“instantiated”) into the model by specifying irreversible binding at these three sites (Fig. 5) to comprise fully specified CaMKII holoenzymes. In BNGL syntax, a bond between two molecules is represented as a period and the components involved in binding are followed by a ! (pronounced “bang”) and a number. For example a bond between the left side of one subunit and the right side of another is represented as CaMKII(l!1, r, c, T286∼0, pp1∼0, T306∼0, cam∼la).CaMKII(l, r!1, c, T286∼0, pp1∼0, T306∼0, cam∼la). The ! can be followed by any number so long as the two binding components have the same number. “T286” is the threonine autophosphorylation site that confers Ca^2+^-independence; it is modeled as having two states, either unphosphorylated (∼0) or phosphorylated (∼P). pp1 is a binding site for the catalytic subunit of protein phosphatase-1; it can have two states, either unbound (∼0) or bound (∼1) to PP1. “T306” is the threonine autophosphorylation site that can be phosphorylated after CaM unbinds from the subunit. The T306 site remained unphosphorylated (∼0) in the present study. The “cam” component is the binding site for calmodulin; it can have two states, either low affinity (∼la), or high affinity (∼ha). The ∼ha state is the high affinity state that appears when a subunit is autophosphorylated at T286; i.e the “CaM-trapping” state (see reaction rules below).

**Figure 5.**
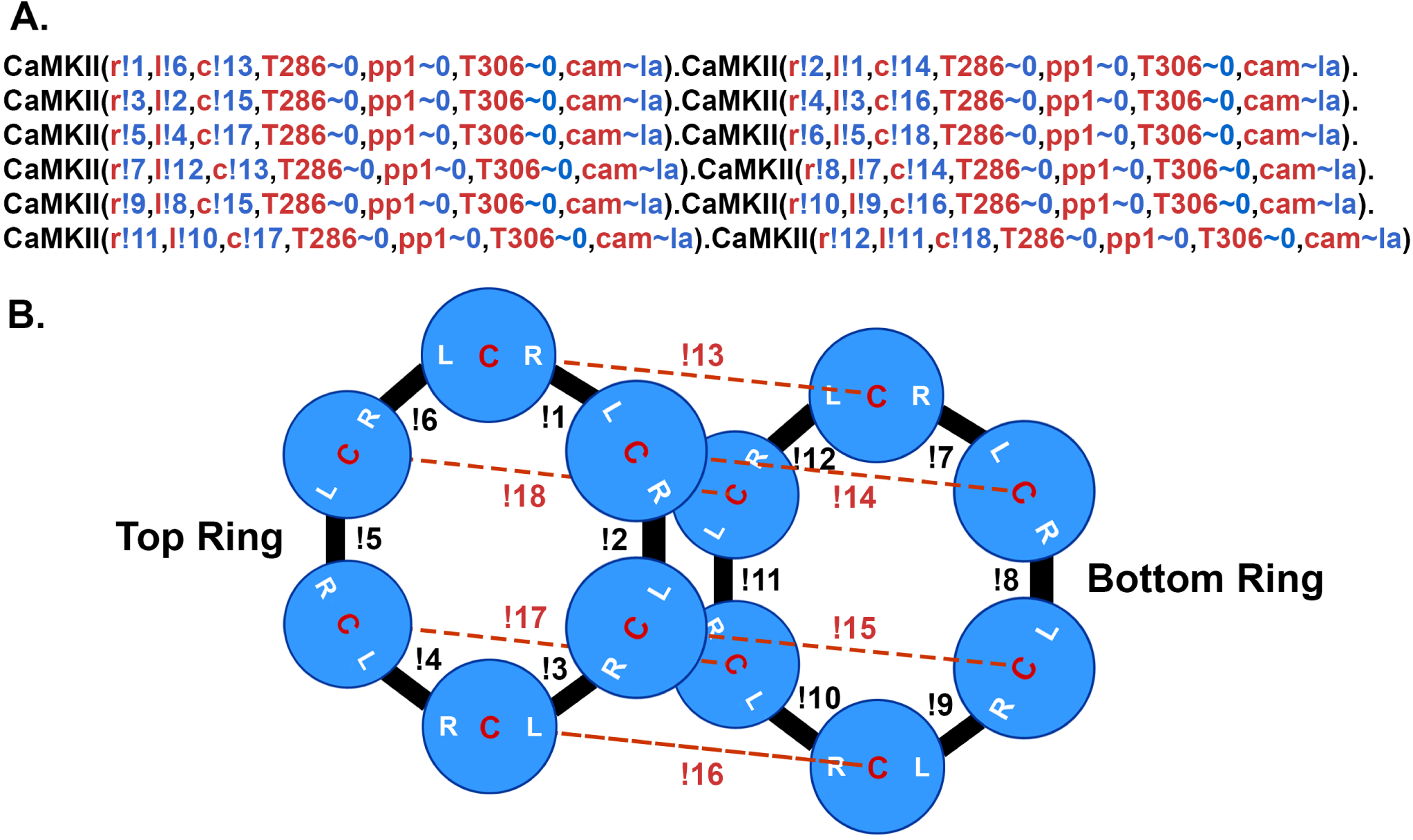
BNGL model of the CaMKII holoenzyme. A. The BNGL code represents a CaMKII holoenzyme as it is initialized in a simulation. A single subunit is defined in BNGL as follows (see Ordyan, et al., 2020): CaMKII(r,l,c,T286∼0∼P, pp1∼0∼1,T306∼0∼P,cam∼la∼ha. Components r, l, and c represent binding sites among subunits in the holoenzyme as diagrammed in B. T286 is a component representing the autophosphorylation site threonine-286 which can be present in one of two states ∼0 (unphosphorylated) or ∼P (phosphorylated). Pp1 is a site for binding of protein phosphatase-1 which can be either unbound (∼0) or bound to PP-1 (∼1). T306 is the autophosphorylation site threonine-306. Cam represents the binding site for Ca^2+^/CaM which is allowed to be in either a low affinity binding-state (∼la) with KD = 65 nM or the high affinity state associated with CaM-trapping (∼ha) with KD = 1.8 pM. Bonded subunits are separated by a period. Bonds between components of subunits are represented by the symbol ! (bang) followed by a number. The number must agree for the two components that are bound to each other. The holoenzyme is initialized with no sites phosphorylated and all cam components in the ∼la state. B. Diagram of the bonds formed among the r, l, and c sites on each subunit to encode the dodecameric holoenzyme.

### Reaction rules

Reaction rules are specified in BNGL as discrete transformations. For example, the rule “CaM(C∼0,camkii) + ca ↔ CaM(C∼1,camkii) k_on1C, k_off1C” encodes reversible binding of one Ca^2+^ to the C-terminus of CaM for CaM not bound to CaMKII with k_on_ = k_on1C and k_off_ = k_off1C. The rate constants for each rule are used to compute the probability that the transformation occurs when the two molecules involved in a bimolecular transformation collide. Unimolecular transformations occur at a future time step obtained by sampling the expected first-order decay time course given by the rate constant. At run time, a random number generator is initialized with a seed number for each individual simulation that specifies a sequence of random numbers that are used to generate the sequence of Brownian motion random-walk trajectories and the probabilities of state transitions for the simulation.

As described above, the model of binding among Ca^2+^, CaM, and CaMKII subunits in the holoenzyme includes rules for binding/unbinding of Ca^2+^ to each of the four sites on CaM when CaM is not bound to CaMKII, binding/unbinding of each state of CaM to a subunit of CaMKII in the holoenzyme, and binding/unbinding of Ca^2+^ to each of the four sites on CaM when CaM is bound to CaMKII. Binding of CaM to CaMKII increases its affinity for Ca^2+^, as expected from thermodynamic detailed balance. These rules and experimental parameters are taken from the model in Pepke et al. (19).

Rules are included for autophosphorylation of one subunit of CaMKII by another. Autophosphorylation at Thr 286 occurs between adjacent subunits in a holoenzyme when they were both active, either because they both have bound Ca^2+^/CaM or because the “right-hand” subunit is already autophosphorylated. We modeled the autophosphorylation reaction as proceeding in one direction around a ring, with the “right-hand” subunit acting as the kinase and the “left-hand” subunit acting as substrate. This situation fits the structural analysis of subunit interactions during autophosphorylation (92). Parameters that govern the probability of autophosphorylation were derived from measurements in Shifman et al. (91). In that study. autophosphorylation rates (turnover numbers) were measured at 30° C with a temperature controlled quench-flow device for CaMKII bound to wild-type CaM4, and bound to mutant CaM’s that bind Ca^2+^ only at the N or C termini (91). We generated 64 permutations of the autophosphorylation rules between neighboring CaMx-bound subunits in a holoenzyme (CaMx denotes CaM with all possible numbers of bound Ca^2+^ ions). Pairs of subunits, both of which contain bound CaM4, undergo autophosphorylation of the left subunit at the rate k_pCaM4 (0.96 s^-1^, Shifman et al., 2006). All other permutations underwent autophosphorylation of the left subunit at the rate k_pCaMpartial (0.1 s^-1^), which is a coarse-grained approximation of the rates measured by Shifman et al. (91). Eight rules are included in which the right-hand subunit of a pair is autophosphorylated but no longer has bound CaM and the left hand subunit contains bound CaMx. For these rules, when the target subunit is bound to CaM4, the rate of autophosphorylation is k_pCaM4. When the target subunit was bound to CaM with less than 4 Ca^2+^ ions bound, the rate of autophosphorylation is k_pCaMpartial.

### CaM-trapping

Meyer and Schulman (23) showed that the initial binding of Ca^2+^/CaM to a subunit within the CaMKII holoenzyme occurs with moderate affinity (K_D_ = ∼50 nM). However, following autophosphorylation, the inhibitory domain is moved entirely away from the substrate pocket (92), leading to a greater than 1000-fold higher affinity for CaM, referred to as CaM-trapping (see Introduction, 23, 62, 63). We added reactions to the model to simulate CaM-trapping in order to enable comparison of simulations with and without CaM-trapping. A component state termed cam∼la was added to CaMKII subunits representing the initial lower affinity CaM-binding site; the component state cam∼ha was added representing the higher affinity site present following autophosphorylation of a subunit. The off rates of individual CaMx species from CaMKII for each state were calculated based on the necessity that ΔG = 0 around a reaction cycle (“detailed balance,” e.g. see 19). We set the rate of transition from cam∼la to cam∼ha following autophosphorylation equal to the rate of dissociation of CaMx from the cam∼la site.

The reactions of CaMKII holoenzymes and CaM and the autophosphorylation reactions were implemented within the spine geometry shown in Fig. 4, which was taken from the model of Ca^2+^ flux described above (12). The inclusion of reactions leading to CaM-trapping by autophosphorylated CaMKII subunits enabled comparison of activation and autophosphorylation kinetics in the presence and absence of CaM-trapping. Inclusion of reactions specifying dephosphorylation by PP1 enabled examination of the effects of variations in the amounts of PP1 activity. Finally, the creation of a version of the model specifying that dephosphorylation by PP1 does not occur when CaM is bound to an autophosphorylated subunit enabled comparison of the kinetics of autophosphorylation with and without blocking of dephosphorylation by bound CaM.

### Postsynaptic EPSPs and Action Potentials

A single 25 mV postsynaptic EPSP resulting from release of glutamate from one vesicle activating both AMPA and NMDA receptors was simulated in the program NEURON. The 25 mV peak was the average size for hippocampal CA1 pyramidal neurons, calculated by Harnett et al. (93). Because the NEURON model did not contain glutamate receptors, the excitatory postsynaptic potential (EPSP) was simulated by injecting current into the spine head as an alpha function such that a 25 mV peak EPSP was produced. The peak of the EPSP occurred about 3 ms after release and decayed with a *τ* of ∼10 ms. The time course of the EPSP was saved to the data file “post_epsp_spine_voltage.dat” to be used at runtime.

The APs in the soma and the associated back-propagating APs (bAPs) in the dendrites arising from the two-epoch stimulus were simulated in the program NEURON by injecting current into the axon hillock of pyramidal neuron, model “j4” (94). The voltages experienced at a spine located on a dendritic branch ∼100 µm from the soma were recorded, as described in Refs. (95) and (12). The spine was chosen so that the bAP arrived at the spine 10 ms after the presynaptic AP. The current injection was large enough to reliably initiate an AP in the soma and subsequent bAP, as it would if applied to an axon bundle with many synapses ending on the neuron. The ion channels included in the j4 simulation are described more fully in Ref. (12) and in the NEURON ModelDB (https://senselab.med.yale.edu/ ModelDB/ShowModel.cshtml?model=2488). The time course of the bAP arriving at the spine was saved to the data file “post_bAP_spine_voltage.dat.”

A python script (vm_dependent_rates_post.py) was written to add together the changes in membrane potential in the spine resulting from EPSPs during simulations for each seed, and from the bAP arriving 10 msecs after the AP in the soma. We assumed that the EPSP and bAP summed together linearly. There was a reliable 10 ms delay between the generation of the presynaptic spike in the soma and the arrival of the postsynaptic bAP at our single spine in simulations for each seed. In contrast, the release of glutamate and the resulting timing of the EPSP were more variable because of the biologically realistic stochastic elements built into the simulation of release (Fig. S1 E). When glutamate was released, it was most often at 10ms before the arrival of the bAP. However, occasionally, for the reasons discussed above, glutamate release and the associated EPSP did not appear causally linked to the bAP. Two such events are shown at ∼2.4 and ∼6.2 sec in Fig. S1 C and E, which depict results of a single simulation for seed #50. NMDARs open and flux Ca^2+^ into the spine only when they bind glutamate at the same time that the EPSP and bAP depolarize the spine. The coincidence of these events opens the channel and relieves the Mg^2+^ block that prevents movement of Ca^2+^ through the channel (83, 96). The complex kinetics of the transitions that allow Ca^2+^ flux through the NMDARs are explained in detail in Ref. (12).

### Data Analysis

The simulations were performed on an HPC cluster at Caltech on 50 nodes each having at least 256 Gbytes of RAM. Monte Carlo simulations in MCell4 (40) rely on generation of random numbers to determine the diffusion trajectories and probabilities of each reaction. Each simulation was initiated with a different seed for the random-number generator. Simulation of 50 seeds for each condition, for 11 seconds of simulated time, required 7 to 10 days of compute time. Output, consisting of numbers of molecular states across time, were stored as ASCII data files. MCell permits output of a “checkpoint file,” that can be used to extend the time of a simulation beyond our standard 11 seconds if desired. The number of individual simulations for each condition required to obtain high confidence averages was determined by comparing the mean and standard deviation of 30, 40, and 50 simulations of calcium dynamics. Our analyses showed that there was no statistically measurable difference between the population averages for 40 and 50 seeds. We thus used 50 seeds for each condition in our simulation to ensure that we would obtain a full representation of all possible stochastic trajectories. All data are plotted as the mean of 50 seeds.

Output was plotted in MatPlotLib (matplotlib.org) or Prism (GraphPad, www.graphpad.com). The time constant (**τ**) of exponential decay of pCaMKII caused by dephosphorylation by PP1 was calculated by fitting the amplitude and decay time of averaged curves of pCaMKII with an equation for a single exponential.

## Materials Availability

The suppementary movie, a readme file, and the eight models used to generate the data reported here are available for download as a compressed file at: http://www.mcell.cnl.salk.edu/models/spatial-model-of-CaMKII-2024-1/. They are also available on Zenodo at https://doi.org/10.5281/zenodo.12764450.

MCell4 is available for download at http://mcell.org/ and as source code at http://github.com/mcellteam/mcell.

## Results

### Calcium dynamics in the spine in response to the two-epoch stimulus

Ca^2+^ influx into the spine through the NMDARs and VDCCs was evaluated in response to the two-epoch stimulus shown in Fig. S1. As in our previous study (12), the model captured the complex, stochastic kinetics of NMDAR and VDCC channel openings (Fig. S2). The NMDAR channel was opened by binding of glutamate, and the Mg^2+^ block of Ca^2+^ flux was relieved by depolarization of the membrane during the bAP (Fig. S2 A, B). During the first epoch, the number of open NMDAR channels decreased for the later three bursts of APs, reflecting the decrease in the number of vesicles releasing glutamate (Fig. S1 C). The bAPs continued to relieve the Mg^2+^ block for all open NMDAR channels (Fig. S2 A, B). The number of open NMDAR channels decreased substantially for the second epoch due to the lower release probability from the presynaptic terminal caused by short term depression (Fig. S1 C) and a small effect of NMDAR desensitization. The stimulus caused voltage-dependent openings of the single VDCC in the spine (Fig. S2 C, D). As expected, the stimulus resulted in calcium influx into the spine through NMDARs and VDCCs (Fig. 6). Fluxes through NMDARs and VDCCs corresponded to the channel opening dynamics shown in Fig. S2. Although the peak Ca^2+^ flux through the VDCC was greater than that through the NMDARs, the integrated flux through NMDARs was considerably greater than the integrated flux through the VDCC, as is evident in Fig. 6B. As a result of these fluxes, the free Ca^2+^ in the spine increased in response to the stimulus (Fig. 6 C, D), as did all of the Ca^2+^-bound species of free CaM (Fig. S3). Total free Ca^2+^ entering during the first epoch was greater than that during the second epoch due to synaptic depression. In this model, CaM can diffuse from the shaft into the spine during the stimulus; thus, the total Ca^2+^-bound CaM and CaM bound to CaMKII exceeded the amount of CaM initially added to the spine alone.

**Figure 6:**
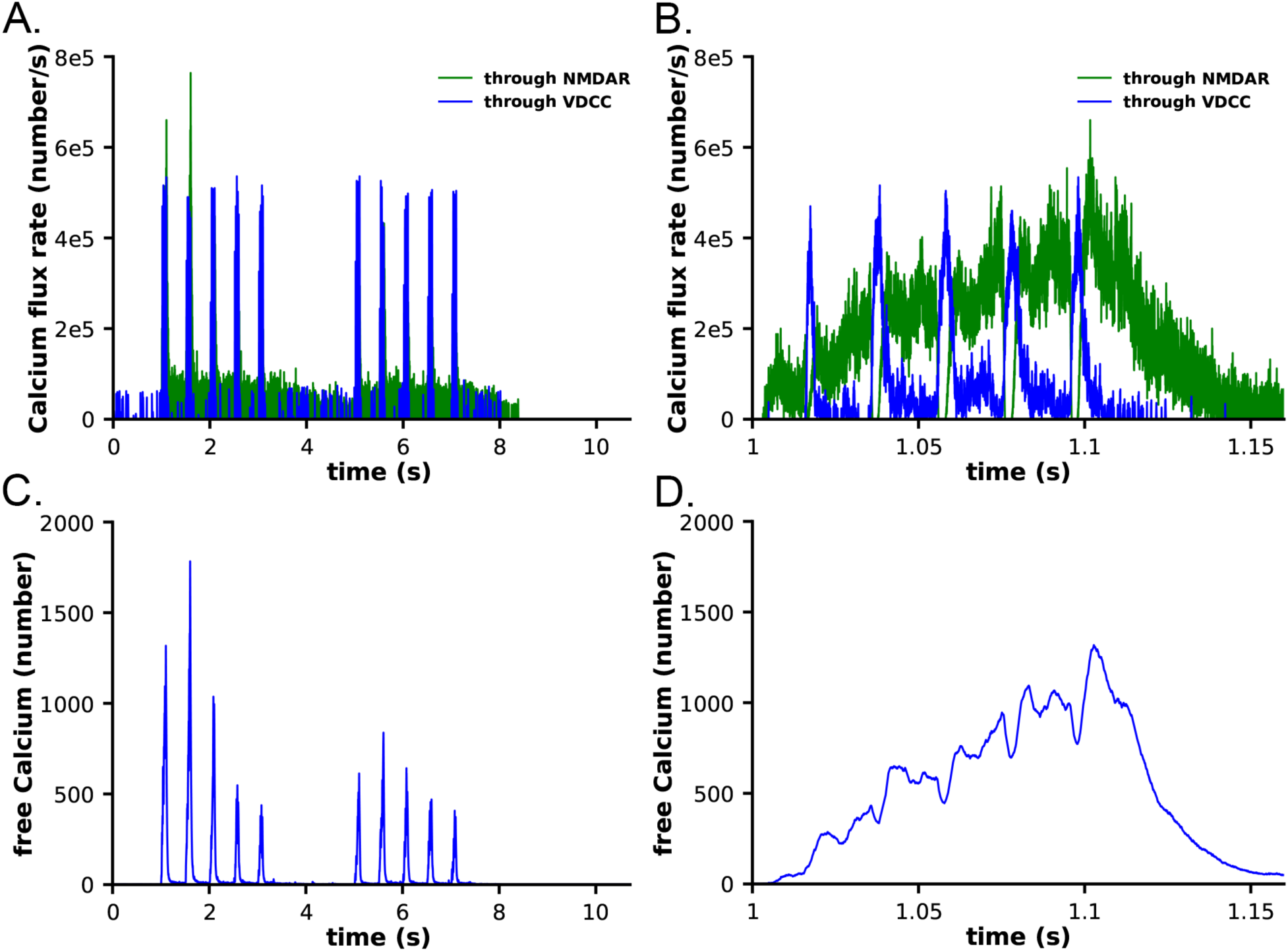
Calcium influx in spines in response to the stimulus shown in Figure 5. **A**. Calcium flux rate through NMDARs (green) and VDCC (blue). **B**. Data from A. between 1 and 1.15 s. **C**. Number of free calcium ions in the spine. **D**. Data from C. between 1 and 1.15 s. Each curve is the average of 50 individual seeds. The dynamics of NMDAR, VDCC, and calcium are the same in the CaM-trapping and non-trapping models. The dynamics of the different CaM species are shown in Fig. S3.

### Sensitivity to protein phosphatase 1 activity

It has been shown experimentally that the dynamics of autophosphorylation of CaMKII and induction of LTP are highly sensitive to the concentration of active phosphatase (57, 97, 98). Most of the simulations in this study were obtained with a uniform concentration of active PP1 (1.25 µM) and of CaMKII (80 µM subunits) throughout the spine. We chose this amount of PP1 as a likely median concentration based on the biochemical measurements of Ingebritsen et al. (87). The PP1 catalytic subunit is regulated and localized *in vivo* by a variety of specialized regulatory subunits (98, 99), including in synapses by neurabin (98, 100) and spinophilin (101). Thus, the level of PP1 activity in synaptic spines is likely to be variable and may be regulated by synaptic activity. To test whether the composite model reproduces the observed sensitivity of autophosphorylation to phosphatase activity, we simulated autophosphorylation of CaMKII, varying the concentration of active PP1 in the spine from 0.65 µM to 5 µM (Fig. 7). The amount of formation of pCaMKII was extremely sensitive to the level of PP1 (Fig. 7). Thus, the composite model reproduces PP1 sensitivity *a priori* within the range of the experimentally measured physiological levels of CaMKII and PPI that we used in the model. It is notable that the numbers and activity of PP1 in the models were not adjusted to confer sensitivity to PP1, rather they were set according to biochemical measurements in the experimental literature. Thus, the model reproduces the sensitivity of induction of LTP to the level of phosphatase activity observed experimentally (43, 97).

**Figure 7:**
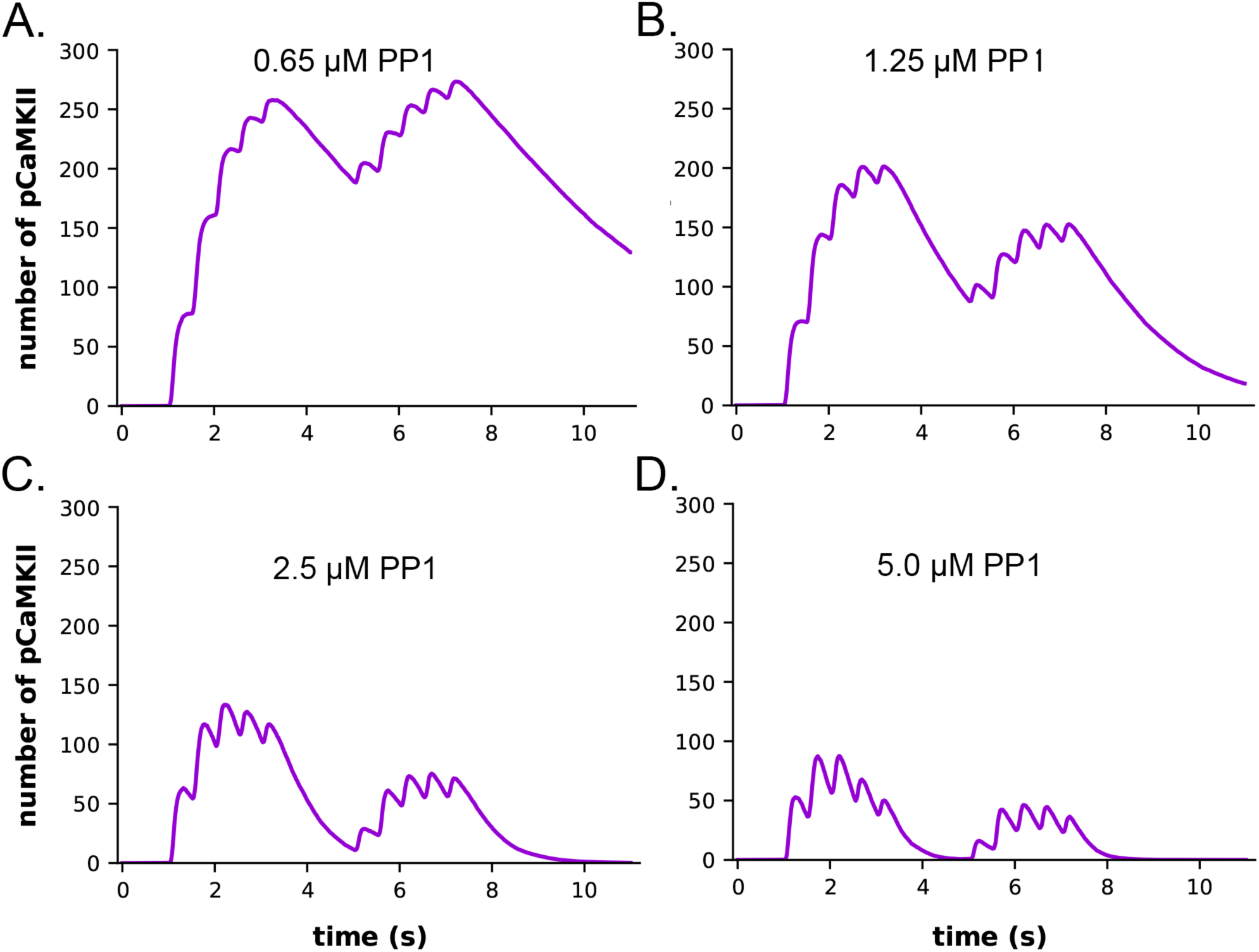
Dynamics of formation of pCaMKII subunits in the presence of increasing concentrations of active PP1 in the CaM-trapping model. **A**. 0.65 μM PP1 **B**. 1.25 μM PP1. **C.** 2.5 μM PP1. **D.** 5.0 μM PP1 Data are essentially identical for the non-trapping model. Each curve is the average of 50 simulations run with different random seeds.

The data also provide quantitative detail about how the interplay between activation or inhibition of PP1 can exert exquisitely tight control of the progression of activation of CaMKII by a given stimulus. When the amount of active PP1 was reduced by half from 1.25 µM to 0.65 µM, the number of autophosphorylated subunits (pCaMKII) remaining at 5 min after the beginning of the stimulus, which was the time of initiation of the second epoch, was more than doubled (from 90 to 192) (Fig. 7). The number of active PP1 molecules bound to a CaMKII subunit at the peak of the first epoch was nearly linearly dependent on the concentration of PP1 between 0.65 and 1.25 µM (Fig. S5). In the lowest concentration of PP1 (0.65 µM), the residual number of pCaMKII remaining after the first epoch (∼190 out of a total of 720 subunits (25%) was sufficient to support an absolute increase in pCaMKII during the second epoch despite the reduced influx of Ca^2+^ (see Fig. 6 C). With 25% of subunits autophosphorylated, the average number of pCaMKII subunits in a six-membered ring would be ∼1.5. In contrast, In 1.25 µM PP1, the average number of pCaMKII was ∼90 out of 720 or ∼12.5%, with an average of ∼0.75 pCaMKII in a six-membered ring. To become newly autophosphorylated, a subunit with newly bound CaMx needs to be located next to an activated subunit in the clockwise direction. A decrease in active PP1 from 1.25 µM to 0.65 µM, increases the probability of this occurring from ∼0.75/5.25 (14%) to ∼1.5/4.5 (33%). This higher probability increases the likelihood that a subunit with newly bound CaMx will be able to be autophosphorylated by its neighbor.

### Dependence of the dynamics of CaM binding to CaMKII on CaM-trapping

We measured the dynamics of formation of phosphorylated CaMKII subunits (pCaMKII) in the absence (Fig. 8A) and presence (Fig. 8B) of CaM-trapping. Each phosphorylated CaMKII subunit could be bound to CaM in any one of eight states; CaM with no bound Ca^2+^ (CaM0), or CaM with 1 to 4 bound Ca^2+^’s (CaM1C, CaM2C, CaM1C1N, CaM2C1N, CaM2N, CaM1C2N, and CaM4). The results show that the total accumulation of pCaMKII was independent of the presence of CaM-trapping with the stimulus applied here. In the presence or absence of CaM-trapping, the numbers of pCaMKII subunits bound to CaM0, CaM1C, CaM1N, CaM2N, CaM1C1N, and CaM1C2N were negligible. In the presence of CaM-trapping, binding of CaM4, CaM2C, and CaM2C1N to pCaMKII was substantially prolonged, and the numbers of these bound CaM species did not fall to zero before the onset of the second epoch (Fig. 8B, D). However, because these species are bound to CaMKII subunits that are already autophosphorylated, the prolonged binding did not increase the likelihood that CaM newly bound during the second epoch would be located next to an active subunit. The presence of trapping did not significantly increase the lifetime of CaM species bound to non-phosphorylated subunits (Fig. S4). For this reason, CaM-trapping did not increase the number of autophosphorylated subunits produced by the stimulus.

**Figure 8:**
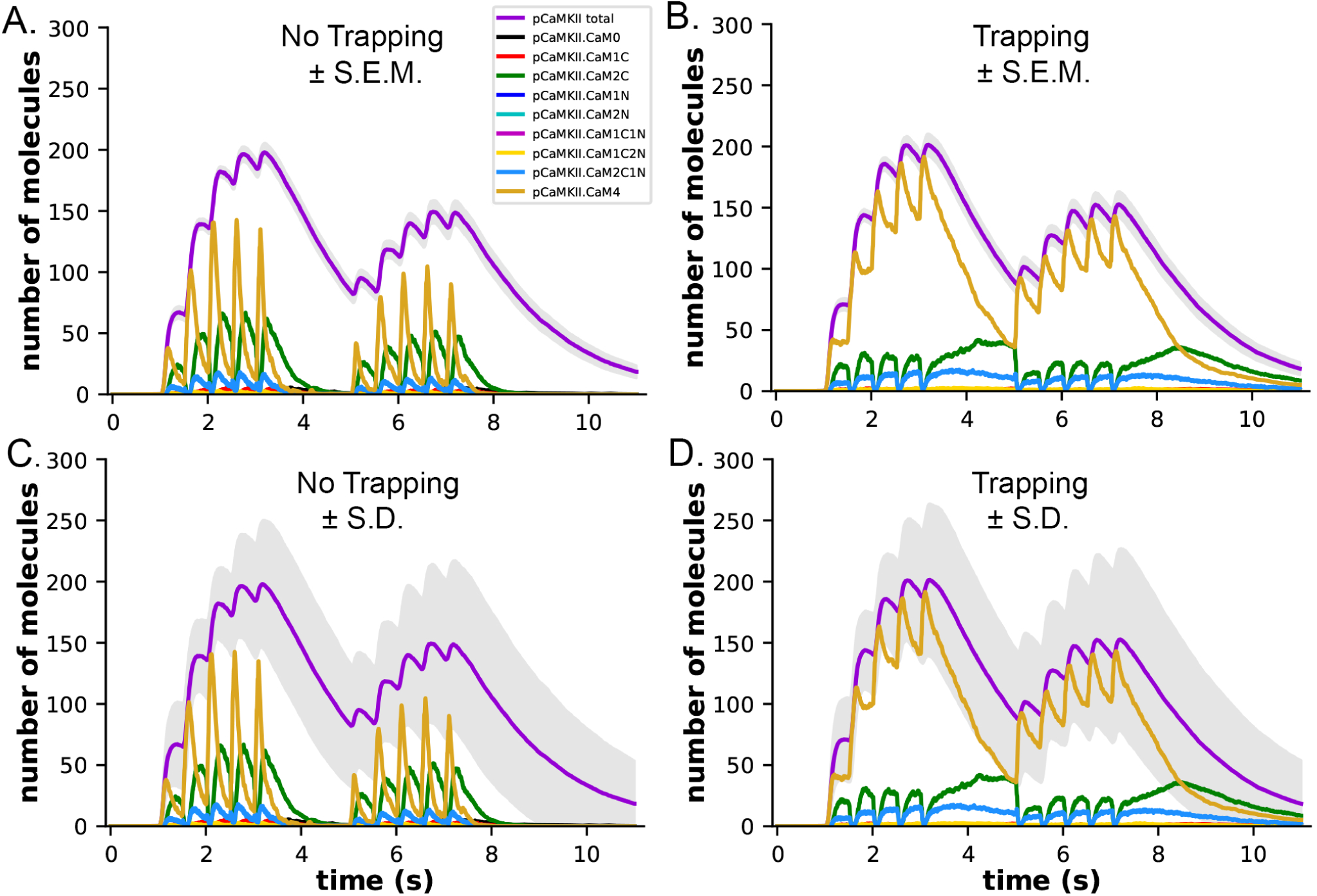
Dynamics of formation of phosphorylated subunits of CaMKII (pCaMKII) and binding of CaM species to them in the presence and absence of CaM-trapping. **A**. Binding of calcium-bound states of CaM to pCaMKII in the absence of CaM-trapping. Colors are as in the legend. **B**. Binding of calcium-bound states of CaM to pCaMKII in the presence of CaM-trapping. Curves are the average of 50 simulations initiated by different seeds. The gray shading surrounding the curve depicting formation of pCaMKII (magenta) indicates +/- s.e.m. **C.** and **D**. The same curves as depicted in A and B, but with the gray shading surrounding the curve depicting formation of pCaMKII indicating +/- s.d. The large standard deviation results from the number of stochastic reactions in the model. The large variation reflects the biological reality of coupled reactions occurring among a limited number of molecules in a small cellular space. See Supplementary Material for a short movie of a simulation with glyphs as shown in Fig. 4 B,C.

Figures 8A and B show the small s.e.m. of formation of pCaMKII averaged over 50 seeds (grey shading), indicating the high accuracy of our calculation of the mean pCaMKII. In contrast, the large s.d. of pCaMKII (Fig. 8C, D, grey shading), captures the large dispersion of data that arose because each seed produces a distinct trajectory of stochastic binding and autophosphorylation. The dispersion is the result of the many probabilistic steps involved in formation of pCaMKII upon Ca^2+^ entry. This “noisiness” reflects the actual physiological variability expected for activation of CaMKII in a median-sized synapse by the stimulus that we used. All simulations results shown are averages of 50 independent simulations run with 50 different random seeds and have similar small s.e.m.’s.

### The effect of localization of CaMKII in the PSD on the dynamics of autophosphorylation

The concentration of CaMKII in the spine cytosol is high; however *in vivo*, a substantial portion of spine CaMKII is further localized to the PSD as shown by immunofluorescence and biochemical studies (102–105), where it is concentrated within ∼50 nm of the postsynaptic membrane. It has been suggested that this portion of CaMKII may be more highly activated and autophosphorylated than cytosolic CaMKII during synaptic stimuli that activate NMDARs (106). To test this idea, we created a capsule associated with the postsynaptic membrane. The capsule has lateral boundaries contiguous with the reconstructed PSD and extends 50 nm into the cytoplasm from the inner membrane leaflet. The volume of the capsule is 0.002915 fL which is ∼ 20% of the spine volume. The walls of the capsule were made reflective to CaMKII and transparent to all other volume molecules. To simulate 50% localization of CaMKII in the PSD, we placed 30 holoenzymes within the PSD capsule (subunit concentration ∼200 µM) and 30 holoenzymes in the spine cytosol outside the PSD capsule (subunit concentration ∼43 µM)., The other variables in this model were identical to those in the simulations shown in Fig. 8 B and D, which included CaM-trapping. Results of simulations with this model are shown in Fig. 9 A and C. Localization of CaMKII in the PSD has only a minimal effect on the proportion of subunits autophosphorylated (Fig. 9C), indicating a small enhancement of autophosphorylation likely arising from the proximity of subunits in the PSD to Ca^2+^ coming through the NMDA receptor. In contrast, as expected, the concentration of pCaMKII in the PSD is approximately 5-fold higher than in the rest of the spine (Fig. 9A). Note that the increased concentration of CaMKII in the PSD does not alter the rate of autophosphorylation upon CaM binding because autophosphorylation is an intraholoenzyme reaction.

**Figure 9.**
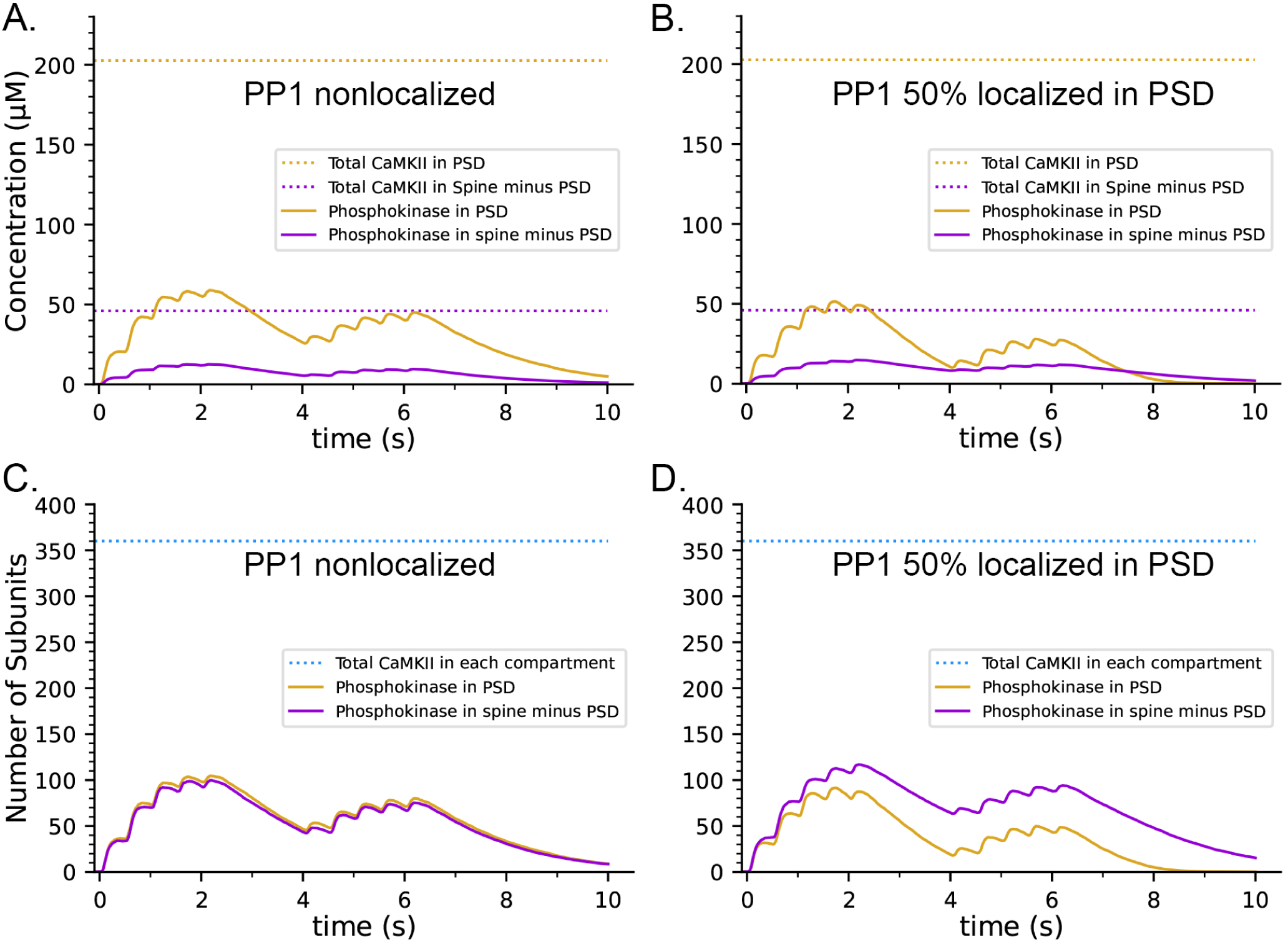
Effects on the dynamics of formation of phosphorylated subunits of CaMKII (pCaMKII) of localization of 50% of CaMKII in the PSD and of 50% of PP1 in the PSD. Lines and colors are as in the legends. **A**. Concentration of pCaMKII in the PSD capsule and in the rest of the spine (minus the PSD capsule) compared to the total CaMKII in each compartment. **B**. Same as in A, but with 50% of PP1 localized in the capsule. **C**. Numbers of pCaMKII in the PSD capsule and in the rest of the spine, compared to the total CaMKII in each compartment. **D**. Same as in C, but with 50% of PP1 localized in the capsule. Curves are averages of 50 simulations initiated with 50 different random seeds.

The higher concentration of CaMKII would, however, enhance the rate of binding of pCaMKII to the nearby GluN2B subunits of the NMDARs (26, 107) and to densin-180 located in the PSD (108, 109), and would increase the rate of autophosphorylation of protein substrates located in the PSD.

It has been reported that pCaMKII is preferentially dephosphorylated by PP1 localized in the PSD (57, 98). To test the effect of localization of PP1 in the PSD, we further modified the model to make the capsule reflective to both CaMKII and PP1; and placed 50% of CaMKII and of PP1 (6 molecules, concentration ∼3.3 µM) inside the capsule leaving the other 50% in the rest of the spine cytosol (concentration ∼0.7 µM). Simulations with this modified model revealed that differential localization of PP1 in the PSD decreased formation of pCaMKII in the PSD and enhanced formation of pCaMKII outside the PSD (Fig. 9 B and D).

### CaM-trapping may influence the rate of dephosphorylation of pCaMKII by prolonging the ability of bound CaM to inhibit binding of PP1

The presence of CaM-trapping did not influence the rate or extent of formation of pCaMKII during the two epoch stimulus that we used here (Compare Fig. 8A and B). However, it did significantly prolong the lifetimes of binding of CaM4, CaM2C, and CaM2C1N to CaMKII (Fig. 8B and D). In an an earlier modeling study (66), it was noted that the CaM-binding domain on CaMKII overlaps significantly with the region on the CaMKII regulatory domain immediately downstream of T286, near where PP1 would bind to catalyze dephosphorylation of T286. Therefore, the authors suggested that bound CaM might block or partially inhibit reversal by PP1 of CaMKII autophosphorylation. The suggestion that bound CaM might interfere with binding of PP1 is supported by structural studies of binding of the PP1 catalytic subunit to substrates (99) and to the targeting subunit spinophilin (110). PP1 interacts with both classes of proteins over a wide area of its surface. This property of PP1 suggests that bound CaM would sterically hinder binding of PP1 to CaMKII in the vicinity of phosphorylated T286, and thus inhibit dephosphorylation.

To measure the effect of CaM-trapping on the postulated inhibition of PP1 dephosphorylation by bound CaM, we modified the reaction rules for dephosphorylation to allow PP1 to bind to a pCaMKII subunit only when the cam site is free. This situation represents the most extreme case in which bound CaM completely blocks PP1 binding. Because a PP1 holoenzyme likely binds to pCaMKII via more than one docking site (99), it is also possible that bound CaM lowers the affinity for PP1 by steric hindrance, but does not block dephosphorylation completely. We did not investigate this latter possibility in the present study.

In both the presence and absence of CaM-trapping, the inclusion of inhibition by CaM of dephosphorylation by PP1 increased the maximum formation of pCaMKII (Compare Fig. 10 A,C to B, D). The combined effect of inhibition by CaM and CaM-trapping was particularly profound, increasing the rate-constant **τ** for dephosphorylation from ∼1.8 s to 60.4 s (Table 1) and resulting in autophosphorylation of approximately half of the CaMKII subunits at peak during the second epoch (Fig. 10D, Fig. 11). Thus, the model predicts that the rate of decay of pCaMKII after a stimulus is intricately coupled to the relative affinities of binding among CaM, PP1, and CaMKII.

**Figure 10.**
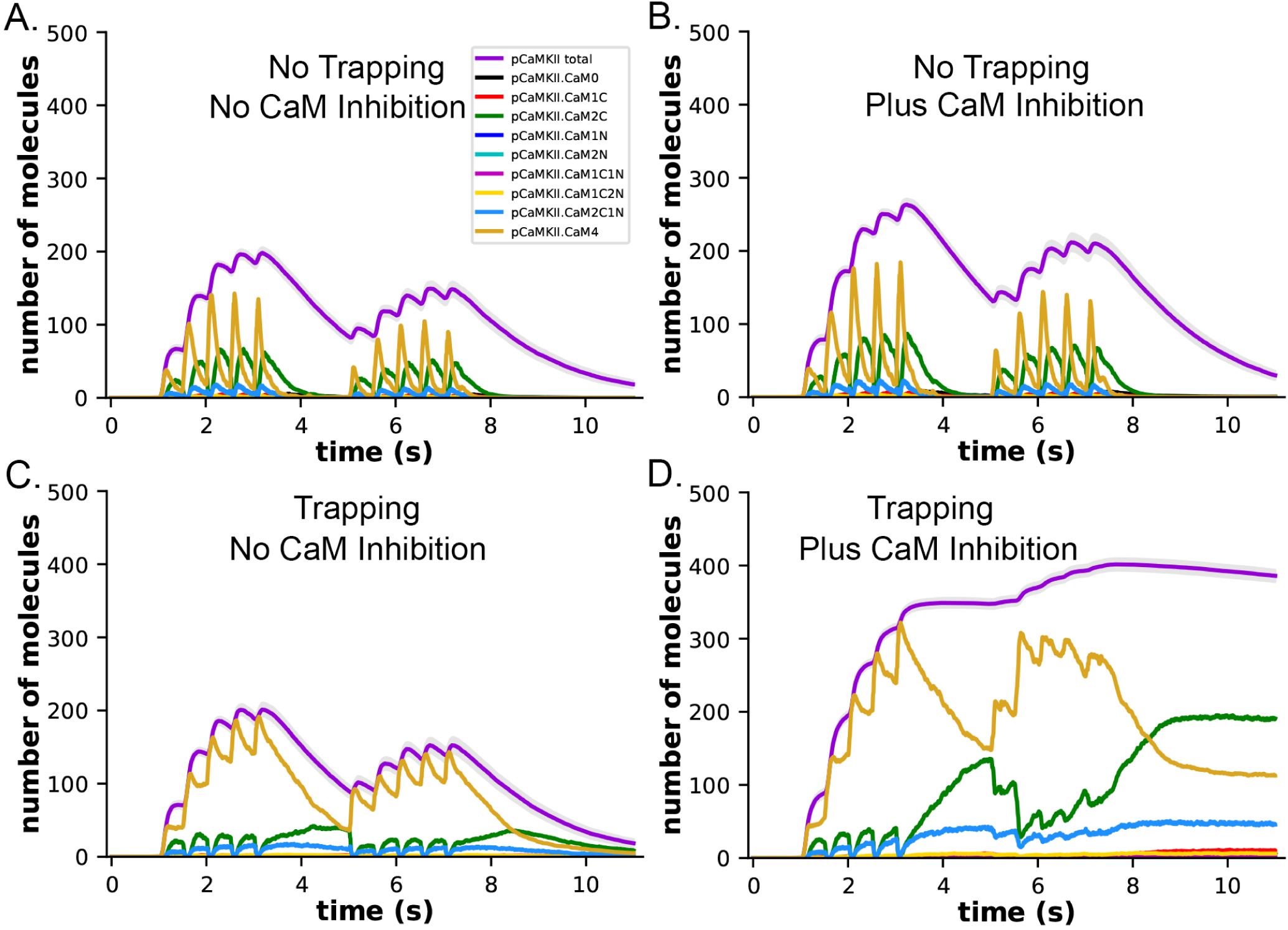
Dynamics of formation of pCaMKII by competition for binding between PP1 and CaM, in the presence and absence of CaM-trapping. **A**. Data replotted from Fig. 8A (no trapping) for comparison to B. **B**. Dynamics of formation of pCaMKII in the presence of competition between CaM and PP1, and in the absence of trapping. **C**. Data replotted from Fig. 8B (trapping) for comparison to D. **D**. Dynamics of formation of pCaMKII in the presence of competition between CaM and PP1, and in the presence of trapping. Each curve is the average of 50 simulations with 1.25 µM PP1, initiated with different random seeds.

**Figure 11.**
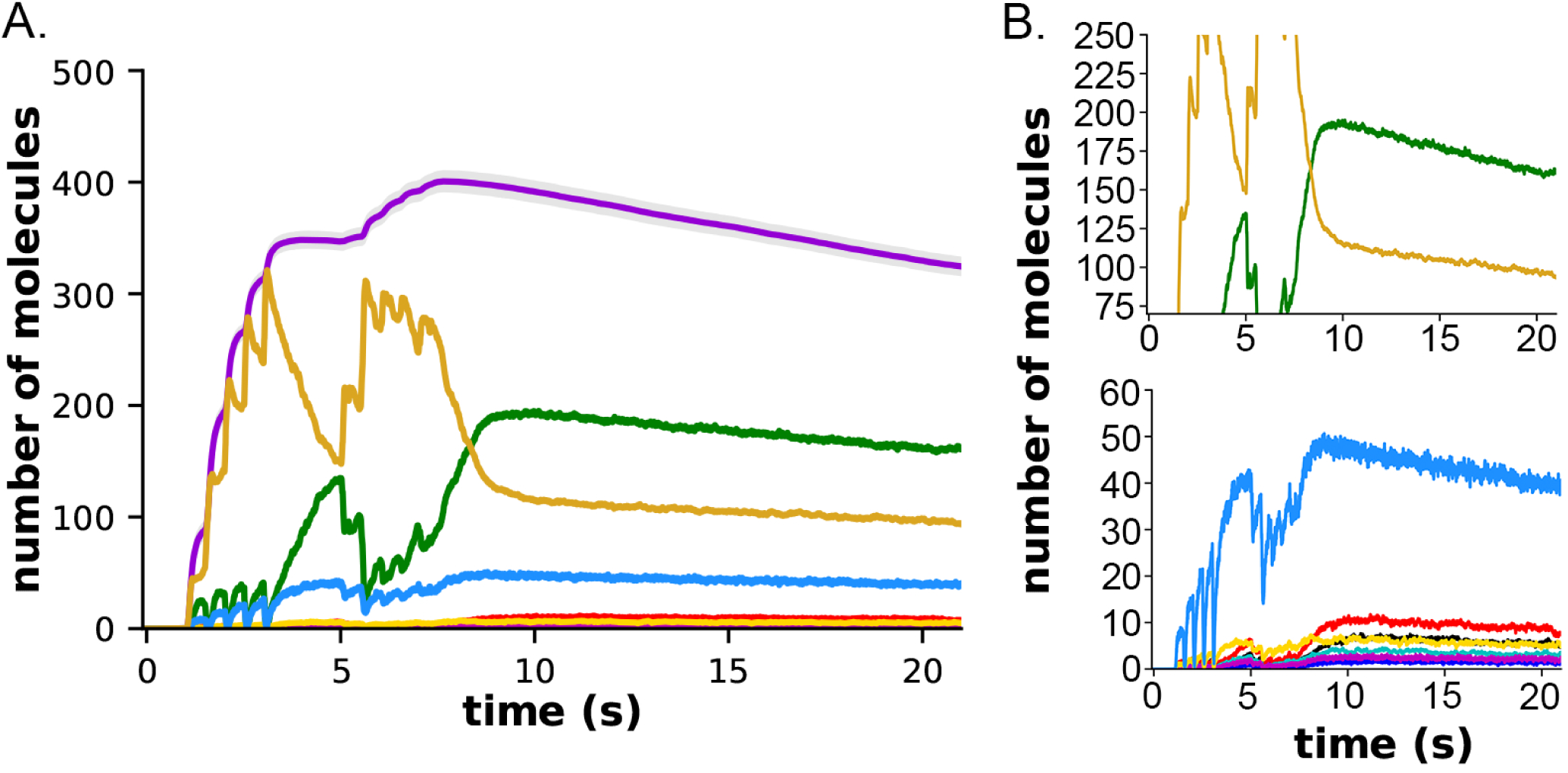
Formation and decay of pCaMKII and bound CaM species after the two-epoch stimulus. **A.** The simulations depicted in Fig. 10D were extended for an additional 10 seconds to measure the rate of dephosphorylation of pCaMKII in the presence of competition for binding between CaM and PP1. **B.** Data from A. plotted with magnified y-axes to show visible decay of bound CaM4, CaM2C (top) and CaM2C1N (bottom). The decay constant for dephosphorylation measured from this data are shown in Table 1.

**Table 1.** Values of tau for dephosphorylation of pCaMKII in the non-trapping and trapping cases were determined at different simulated concentrations of PP1 from the data shown in Figure 7, 8 and data-not-shown for the non-trapping case, as described under Methods. The values of tau in the presence of competition for binding between CaM and PP1 were determined for the non-trapping case from the data shown in Fig. 10 B, and for the trapping case from Fig. 11. The tau for Epoch 1 was identical to that of Epoch 2 as determined by measuring the decay of pCaMKII from the peak at 4 secs to 5 secs, and for Epoch 2 from the peak at 7.7 secs to 8.7 secs. Tau’s for CaM4, CaM2C, and CaM2C1N from Fig. 11B were 61.7, 57.2, and 59.2, respectively.

| [PP1] | $\tau$<br>Non-trapping | | $\tau$<br>Trapping | | $\tau$<br>Non-trapping<br>CaM vs. PP1<br>competition | | $\tau$<br>Trapping<br>CaM vs. PP1<br>competition | |
| --- | --- | --- | --- | --- | --- | --- | --- | --- |
|  | Epoch 1 | Epoch 2 | Epoch 1 | Epoch 2 | Epoch 1 | Epoch 2 | Epoch 1 | Epoch 2 |
| 0.65 μM | 5.0 s | 4.9 s | 5.01 s | 4.8 s | | | | |
| 1.25 μM | 1.9 s | 1.7 s | 2.0 s | 1.8 s | 2.3 s | 1.9 s | 60.4 s | 60.4 s |
| 2.5 μM | 0.74 s | 0.65 s | 0.74 s | 0.67 s | | | | |
| 5.0 μM | 0.33 s | 0.32 s | 0.38 s | 0.32 s | | | | |

To evaluate the effect of inhibition by CaM when both CaMKII and PP1 are 50% localized in the PSD capsule, we added the inhibition rules to the localization model shown in Fig. 9. In simulations with the modified model, the maximum concentration of pCaMKII formed in the PSD during the stimulus increased almost 3-fold from about ∼50 µM (Fig. 9B) to ∼140 µM (Fig. 12 A). Although ∼67% of subunits were autophosphorylated at the peak, the concentration of pCaMKII continued to decay slowly when the stimulus ended (Fig. 12 A, B). With 50% of CaMKII and PP1 both sequestered in the PSD capsule (Fig. 12), the rate of decay (**τ**) in the capsule was 54.5 s and the rate in the cytosol outside the capsule was 72.0 s, compared to 60.4 s with neither sequestered (Table 1). These results mean that, under these physiological conditions, dephosphorylation by PP1 is not overwhelmed by the rate of recurring autophosphorylation within the holoenzyme, even when maximum autophosphorylation reaches 67% of the total subunits.

**Figure 12.**
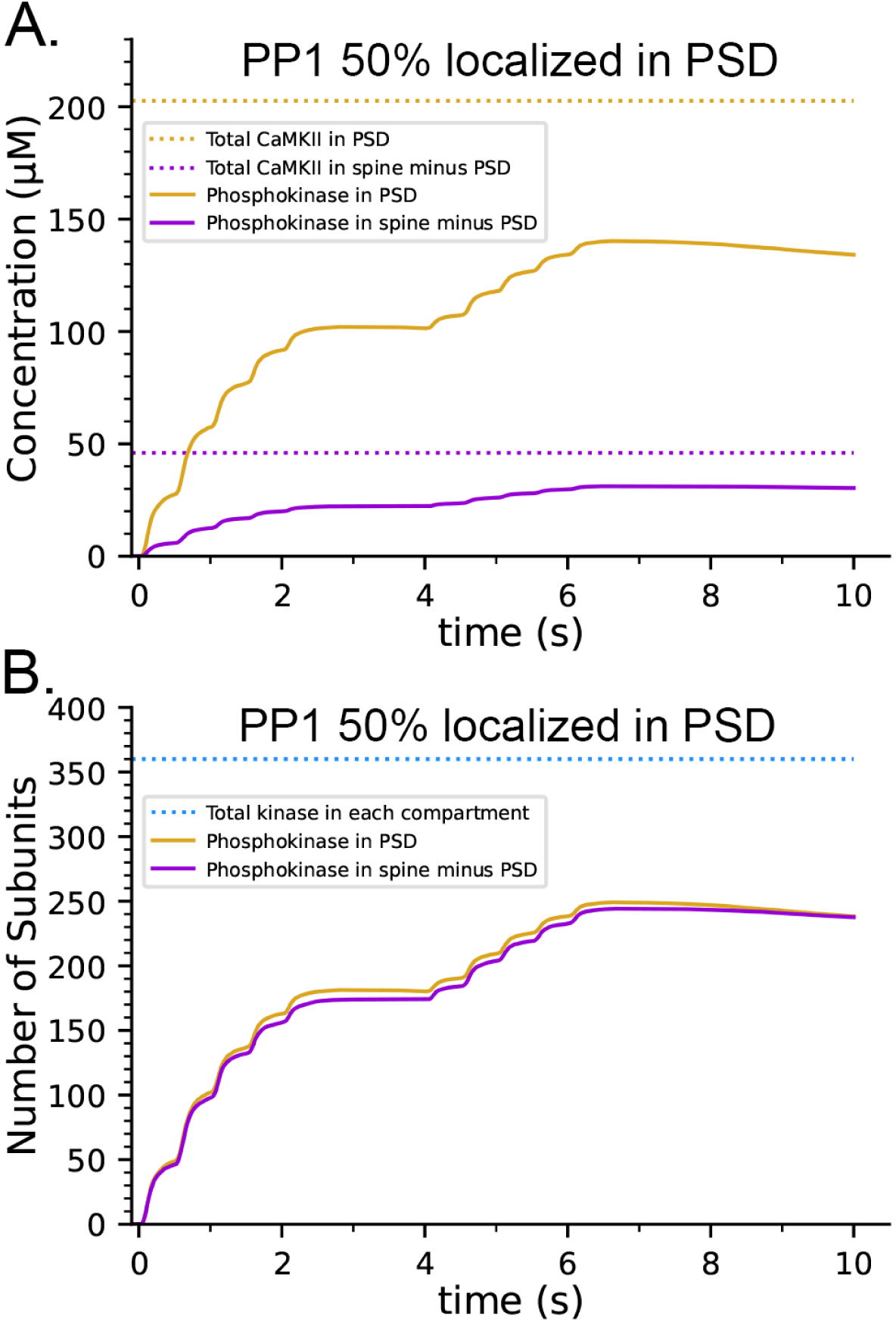
Effect of inhibition of PP1 activity by bound CaM on autophosphorylation when both CaMKII and PP1 are differentially localized in the PSD. **A**. The concentration of pCaMKII reaches ∼67% of the possible maximum of ∼200 µm in the PSD. **B**. The proportion of autophosphorylated subunits in a holoenzyme is ∼67% in both compartments. Curves are averages of 50 simulations initiated with 50 different random seeds.

## Discussion

We developed a series of agent-based stochastic computational models to examine the predicted responses of CaMKII holoenzymes during the first 20 secs after a two-epoch stimulus of medium strength in a realistic median-sized spine. The models are based upon four earlier carefully-vetted published models (12, 19, 68, 70). They contain physiologically realistic numbers of proteins situated in a spine from hippocampal neuropil (12). The numbers and kinetic parameters of the added CaMKII holoenzymes, CaM, and PP1 are drawn from measurements made over several years during biochemical and electrophysiological experiments cited in the text. Parameters are adjusted to 30-35° C. The use of experimentally measured parameters and realistic numbers and spatial arrangements of the relevant proteins enabled testing of several hypotheses about the behavior of CaMKII under conditions present *in vivo*, that are difficult or impossible to test directly with experiments.

Previous experimental evidence has shown that PP1 is specifically enriched in synaptic spines (57, 111), and regulates the direction and magnitude of changes in synaptic strength produced by different frequencies of stimulation (98). For example, the relationship between stimulus intensity and the direction of change in synaptic strength is altered by pharmacological agents that inhibit PP1 phosphatase activity (43, 112). Under baseline conditions in slices from hippocampal area CA1, low frequency stimulation (1-3 Hz) leads to induction of long-term depression (LTD); stimulation at ∼10 Hz is “neutral,” producing no change in synaptic strength, and higher frequency stimulation in the range of 25 -100 Hz induces long-term potentiation (LTP) (43, 113). However, induction of LTD by stimulation at 1 Hz is blocked by application of inhibitors of PP1 (114). Conversely, induction of LTP is potentiated by inhibition of PP1 (115). Cyclic AMP-regulated inhibition of PP1 acts as a “gate” that determines whether LTP is produced by a given stimulus. Activation of the cAMP pathway by a variety of pharmacological agents permits induction of LTP by both widely spaced “HFS” stimulation (3 pulses of 900 APs at 100 Hz separated by 10 min, 97) and by “θ pulse” stimulation in the 5-12 Hz range (43).

When the increase in cAMP is blocked, these stimuli do not induce LTP. Activation of β-adrenergic receptors by noradrenaline also increases LTP at hippocampal synapses through the cAMP pathway that inhibits PP1, suggesting that a similar mechanism underlies enhancement of learning by noradrenaline (116). We chose the concentration of PP1, and its enzymatic parameters according to what we considered the best estimates of these parameters from the biochemical literature. Thus, the agreement between experimental findings showing sensitivity of synaptic plasticity to PP1 activity and the sensitivity of autophosphorylation to PP1 activity in our simulations (Fig. 7), is an important first validation of the combined model.

CaM-trapping is a phenomenon discovered by Meyer and Schulman (23) and elaborated on by Waxham (62). We decided to test the hypothesis that CaM-trapping causes an enhanced, non-linear increase in autophosphorylation of CaMKII during high frequency synaptic stimulation as suggested in earlier publications (9, 64, 65). To do this, simulations were performed in the presence and absence of the reaction rules that specify the change in affinity for CaM resulting from CaM-trapping. The simulations show that CaM-trapping does indeed prolong the association of CaM with autophosphorylated CaMKII subunits, but it does not alter the extent of autophosphorylation during the repeated stimuli in the 50 Hz frequency range that we used here. This is because the association of CaM with non-autophosphorylated subunits is not prolonged by trapping, and, thus, the probability of autophosphorylation is not increased during the second epoch.

We did, however, find that a function of CaM-trapping may be to dramatically alter the rate of reversal of autophosphorylation by dephosphorylation after a stimulus has ended. This finding is based on the structural prediction that binding of CaM to an autophosphorylated subunit of CaMKII is likely to sterically hinder binding of PP1 required for dephosphorylation of T286 (66). The PP1 catalytic subunit interacts with substrates over an unusually large surface area comprising three different surface domains that surround the active site (99). The recognition pockets for PP1 on substrates are usually more distant from the phosphorylated residue than the typical short consensus recognition sequences for protein kinases (98). Binding of PP1 to a substrate via two or more of its surface domains provides avidity that increases the K_M_ and thus, the rate of dephosphorylation. Conversely, reducing interaction of a substrate with one of the surface domains reduces the K_M_ and slows the rate of dephosphorylation (99). In dendritic spines, the predominant targeting subunit for PP1 is spinophilin (101, 117), which binds to spine actin filaments (118). The spinophilin-PP1 holoenzyme, with a molecular mass of ∼150 kDal, is much larger than a CaMKII subunit. Bound spinophilin obstructs one of the three substrate-binding patches on the surface of the PP1 catalytic subunit (110). Thus, the spinophilin-PP1 holoenzyme would be expected to associate with substrates, including CaMKII, via the two remaining binding patches near the PP1 active site (99). CaM binds to CaMKII at a site located ∼ 7 residues downstream of T286 (Fig. 2). Therefore, it is likely to partially, or fully, sterically hinder access of PP1 to phosphorylated T286.

To investigate the effect of competition between CaM-binding and dephosphorylation by PP1, we simulated the most extreme possibility which is complete block of dephosphorylation of T286 by bound CaM by adding a rule that PP1 cannot bind to pCaMKII when CaM is bound. In the absence of CaM-trapping, blocking of PP1 by bound CaM increased the rate of dephosphorylation (**τ**) by ∼10-20% (Table 1). In contrast, when CaM-trapping was included in this model, dephosphorylation was dramatically slowed (Fig. 13;). The **τ** for dephosphorylation was increased 30-fold to 60.2 sec, reflecting the much longer k_off_ for trapped CaM. When bound CaM inhibited dephosphorylation of pCaMKII, the number of autophosphorylated subunits per CaMKII holoenzyme at the peak of the two-epoch 50 Hz stimulus doubled from ∼200 pCaMKII subunits to ∼400, out of the total of 720 subunits. Even this extreme competitive effect would prolong the lifetimes of autophosphorylated subunits for only a few minutes after the Ca^2+^ concentration returns to baseline. The slower dephosphorylation means that some pCaMKII subunits would remain phosphorylated for as long as 4 minutes (4**τ**). This finding is consistent with the experimental observation of Yasuda et al. (44) that increased CaMKII phosphorylation is detectable in dendritic spines for a few minutes after two-photon glutamate uncaging to activate NMDA receptors. Thus, it also constitutes a second validation of our composite model against experimental results.

**Figure 13.**
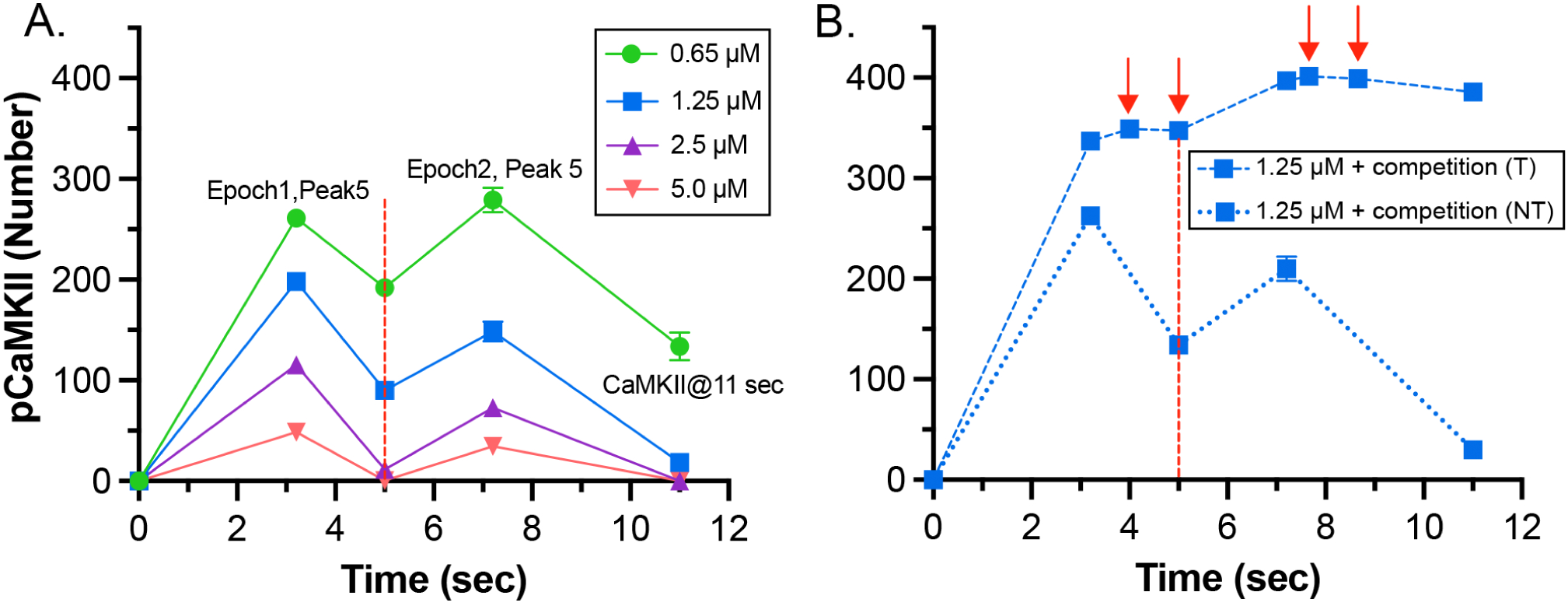
A. Peak concentrations of pCaMKII during the two epoch stimulus in the presence of increasing concentrations of active PP1. Data are replotted for comparison from the results shown in Fig. 7. The dotted red line at 5 sec marks the time of initiation of the second epoch of the stimulus. Averages ± s.e.m. of 50 simulations. **B**. Peak concentrations of pCaMKII during the two epoch stimulus including competition between binding of CaMx and PP1. Data with (T) and without CaM-trapping (NT) are replotted from the results shown in Fig. 10 B and D. In the graph of the model that includes CaM-trapping, points at 4 and 5 sec and at 7.7 and 8.7 secs are marked with red arrows. The differences in numbers of pCaMKII at these times were used to compare the decay time after the first and second epochs. S.e.m.’s for most of the measurements are smaller than the symbols.

Competition between CaM-binding to pCaMKII and dephosphorylation of T286 by PP1 has not been adequately investigated experimentally. The actual effect of CaM-trapping *in vivo* on the rate of dephosphorylation of T286 by PP1 may be less than 100%. The only experimental study we are aware of that indirectly addressed the competition is Bradshaw et al. (88). These authors sought to provide evidence for a bistable kinase switch comprised of CaMKII and PP1. Their experiments indicated that bound CaM did not interfere with dephosphorylation of Thr286 by PP1. However, the experiments were carried out at 0° C to suppress autophosphorylation of Thr305/306; in addition, they used a ratio of PP1 to CaMKII of 1/10, considerably higher than would be predicted *in vivo* from the measurements that we used in our model (∼1/80). We propose that *in vitro* experiments with purified proteins to measure the rate of dephosphorylation of autophosphorylated CaMKII by PP1 in the presence and absence of Ca^2+^/ CaM would be an important test of the effect of CaM-trapping on the persistence of pCaMKII after a strong synaptic stimulus.

Many computational models of activation of CaMKII, incorporating various levels of detail and complexity, have been created previously. One conceptually attractive idea has been particularly persistent in these studies — the notion that CaMKII and PP1 together may form a bistable switch that shifts between an active highly autophosphorylated stable steady-state reflecting LTP and an inactive non-autophosphorylated stable steady-state reflecting LTD (37, 106, 119–123). Because the notion of bistability focuses on steady state behavior, it has been encoded in deterministic equations that specify multiple steady states (121). These models usually include a control parameter that can change the steady state from the low stable state to the high stable state and have been used in mathematical biology to hypothesize that this behavior can occur in complex biological systems (For an introduction, see 124.). As a result, the notion of bistability of CaMKII activity, while appealing in its simplicity, did not evolve from the measured biochemical interactions of CaMKII, CaM, protein phosphatases, or other proposed key proteins. The composite model that we present here is distinct from these earlier models which employed deterministic methods and did not attempt to deduce kinetic mechanisms involving small numbers of molecules and highly structured reaction spaces. Here we have examined the stochastic behavior of CaMKII that is dictated by our best experimental understanding of the numbers of proteins, rate constants of interaction, and subcellular structure in which CaMKII is activated in the first minutes of a complex synaptic stimulus (12, 19, 68, 70). With our models, and other previous models that incorporate more biochemical detail (66, 125), bistability of CaMKII modulated by PP1, as envisioned in Ref. (120), is not observed.

We show that the presence of CaM-trapping in combination with inhibition by CaM of PP1 binding to pCaMKII can lead to a prolonged, but not permanent, elevated level of pCaMKII following the end of a stimulus, perhaps lasting several minutes. This finding suggests that a form of kinetic regulation may be operating that is sometimes referred to as “kinetic proof-reading” (126, 127). This mechanism involves a biochemical network that is structured in such a way that a series of reversible biochemical steps can lead to an essentially irreversible one. The irreversible step only occurs when the preceding reversible steps (for example, binding between two molecules) last long enough to trigger the next step. The canonical example of kinetic proof-reading is the selection of the correct amino acid-tRNA complex to match a codon on ribosomal bound mRNA during protein synthesis. Only an exact match between the tRNA and mRNA codons results in binding that lasts long enough for irreversible formation of the peptide bond to occur (126).

We propose that to induce LTP, autophosphorylation of CaMKII subunits does not need to reach a the high, irreversible steady-state postulated by the bistable switch theory (e.g. 120, 128). Rather, the convergence of signals at the synapse must simply result in a transient high level of autophosphorylation that lasts long enough to trigger one or more downstream, essentially irreversible, steps causing potentiation of the synapse. One such ‘irreversible’ step might include binding of a critical number of active CaMKII subunits to the carboxyl tails of GluN2B subunits of NMDA-receptors (27, 107, 129).

Another might be sufficient dissolution of the protein condensate between PSD-95 and synGAP to trigger irreversible remodeling of the PSD by addition of AMPARs and enlargement of the supporting actin cytoskeleton (5, 29, 72, 130). Both of these events have been proposed to be critical for development and maintenance of LTP.

Several distinct signals can converge on a synapse to modulate the extent of autophosphorylation of CaMKII. The primary signal observed experimentally is repeated high frequency (50-100 Hz) synaptic activation that causes firing of the neuron (see 9, 17). In addition, activation of the cAMP pathway can inhibit PP1, prolonging the lifetime of autophosphorylated CaMKII (32, 43, 97, 116, 131). Src and Fyn protein kinases, activated by G-protein-coupled receptors, receptor tyrosine kinases, and/or cytokines increase the flux of Ca^2+^ through NMDARs, potentiating induction of LTP by high frequency stimulation (132). Kinetic proof-reading would allow combinations of these signals that lead to sufficiently prolonged autophosphorylation of CaMKII in a synapse to induce LTP.

The version of kinetic proof-reading that we describe here is similar, but not identical, to the more abstract “cascade” type models proposed by theorists (35, 133). It attempts to describe how convergence, within a few seconds or minutes, of different experimentally-measured, reversible biochemical signals at a synapse might trigger a structural change that would produce a relatively long-lasting increase in synaptic strength. It remains to be seen whether the kinetic mechanism we propose is related to the “cascades” proposed by theorists.

The MCell4 model presented here can be expanded to investigate the effects of additional proteins and spatial constraints on regulation of CaMKII and the downstream effects of autophosphorylation. Future investigations will include responses of CaMKII in spines of different sizes, the quantitative effect on autophosphorylation of more prolonged and/or lower and higher frequency stimuli.

Importantly, we will investigate the effects of proteins that localize CaM and compete for binding of CaM, and of proteins that regulate PP1. MCell4 is designed to be a powerful tool for introduction of each of these elements in a stepwise fashion to permit dissection of the effects of each and ultimately an understanding of their combined effects.

## Acknowledgments

We acknowledge Prof. T. J. Sejnowski for funding support for T.M.B and M.O, the Air Force Office of Scientific Research (AFOSR) Multidisciplinary University Research Initiative (MURI) grant FA9550-18-1-0051 to P.R., NIH grants MH115456 to M.B.K., DA030749 and MH129066 (CR-CNS) to T.J.S. and M.B.K., and NSF NeuroNex DBI-1707356, NSF NeuroNex DBI-2014862, and NIH MMBioS P41-GM103712 to T.J.S and T.M.B.

## Supplementary Material

Link to supplementary CaMKII_multiburst_color.mov file and legend; and to .tar.gz file containing the four model files (.blend) used in this work: http://www.mcell.cnl.salk.edu/models/spatial-model-of-CaMKII-2024-1/

Legend for “CaMKII_multiburst_color.mov”:

The movie shows the activation of CaMKII holoenzyme within the cytoplasm of a dendritic spine between t=0.9 sec and t=2.1 sec, illustrated in Fig 3 in response to the stimulus shown in Fig 5. The stimulus begins at t=1.0 sec. The model shown in the movie includes all the molecules, reaction pathways, and resulting dynamics described in the manuscript. However, for clarity the movie visualizes only the freely diffusing Ca2+ ions (gold spheres), diffusing molecules of CaMKII holoenzyme (twin torus and sphere glyphs, with color scheme given below), and a patch of 15 NMDARs in the postsynaptic membrane (pentagonal receptor glyphs, with color scheme given below).

Changes in the state of the CaMKII holoenzyme and NMDARs are indicated by changes in color:

NMDAR

unbound: dark green

single Glu bound: medium green

double bound closed: bright green double bound open, unblocked by

Mg: white double bound open, blocked by Mg: red desensitized: dark gray

CaMKII

CaMKII is a dodecamer composed of two hexameric rings (Fig 1). The six subunits of each ring can be autophosphorylated when activated. Here we represent each ring of the holoenzyme by a 2 piece glyph composed of a torus (whose color indicates the CaM-bound state of the hexameric ring) with a sphere at the center (whose color indicates the phosphorylation status of the whole ring). The dodecameric holoenzyme is represented by two of these torus-with-sphere assemblies stacked back-to-back. The color scheme is:

Torus representing a single hexameric ring:

0 CaM4 bound: red

1 CaM4 bound: green

2 CaM4 bound: blue

3 CaM4 bound: cyan

4 CaM4 bound: magenta

5 CaM4 bound: yellow

6 CaM4 bound: white

Sphere representing the phosphorylation status of a single hexameric ring: 0 subunits phosphorylated: black

1 subunits phosphorylated: red

2 subunits phosphorylated: green

3 subunits phosphorylated: blue

4 subunits phosphorylated: cyan

5 subunits phosphorylated: magenta

6 subunits phosphorylated: yellow

## Supplementary Methods

### Numbers of protein molecules

#### CaMKII

The number of subunits of CaMKII in the median-sized hippocampal spine (#37) was set as follows. The average weight of a rat brain is ∼ 1.4 g. A common assumption is that protein makes up about 10% of that weight, or 140 mg. Since ∼80% of the volume of the brain is water and 1g of water has a volume of 1 ml, we assume that the average volume of a rat brain is 1.4 mls. Then the average protein concentration in the brain is ∼140 mg/1.4 ml = 100 mg/ml. We found that CaMKII is highly concentrated in the brain (1). In the hippocampus it is ∼2% of the total protein (2). Thus, its concentration by weight is ∼ 2 mg/ml, averaged over hippocampal tissue. Since it is found almost entirely in neurons, we assume that its average concentration in hippocampal neurons is ∼ 4 mg/ml. The average concentration of the ∼600,000 kDal holoenzyme in hippocampal neurons is:

4 mg/ml x 1 mmole/600,000 mg x 1000ml/liter = 0.00667 mM or 6.67 µM. The average concentration of individual CaMKII subunits is 12 x 6.67 = ∼80 µM.

The volume of the cytosol of spine #37 is 0.016 µm^3^ (0.016 fl). So, the number of holoenzymes in the spine cytosol is:

∼ 1.6 x 10-17 liters x 6.67 x 10-6 (moles/liter) x 6.02 x 1023 (particles/mole) ≈ 64.

We set the number of CaMKII holoenzymes in the spine to 60; and the number of individual CaMKII subunits to 720 (∼80 µM).

#### Calmodulin

The concentrations of CaM in soluble and particulate fractions of brain were measured by Kakiuchi et al., 1982 (3). Estimates of the total concentration ranged from ∼25 to ∼35 µM. We added 30 µM CaM to the spine and to the attached cylinder. The initial number of CaM molecules added to the spine was 290. Because CaM was allowed to diffuse freely between the spine and the attached cylinder, the total number of CaM molecules (bound and free) in the spine increased during the stimulus as CaM bound to CaMKII and free CaM stayed constant. See Figs. 7S and 8.

#### Protein Phosphatase-1

The absolute concentration of PP1 in hippocampal spines is not well measured experimentally. We first estimated an overall concentration in brain from data in References (4) and (5). We then took into account the findings of Shields et al. (6), as well as Allen et al. (7) and Ouimet et al. (8), all of whom show that PP1 is highly concentrated in synaptic spines compared to overall brain cytosol.

Ingebritsen et al. (4) measured the overall specific activity of PP1 in brain tissue as 0.58 nmoles Pi/min/ mg protein. If 1 liter of brain volume contains 100 g protein, we get an approximate average concentration of PP1 activity in the brain overall = 58,000 nmoles Pi/min/liter.

Watanabe et al. (5) measure the k_cat of PP1 as:

∼11.5 Pi released /s/PP1 molecule = 690 Pi/min/PP1 molecule = 690 nmol Pi/min/nmol PP1. Thus, we can calculate an approximate concentration of PP1 catalytic units in the brain overall as: 58,000 nmoles Pi/min/liter x 1 nmole PP1/690 nmol Pi/min = 84 nmoles/liter = 0.084 µM.

Because PP1 is considerably more concentrated in spines than in brain overall, we used a concentration of 1.25 µM as a starting point. We also simulated the effects of 0.65 µM, 2.5 µM and 5 µM.

**Table S1:**
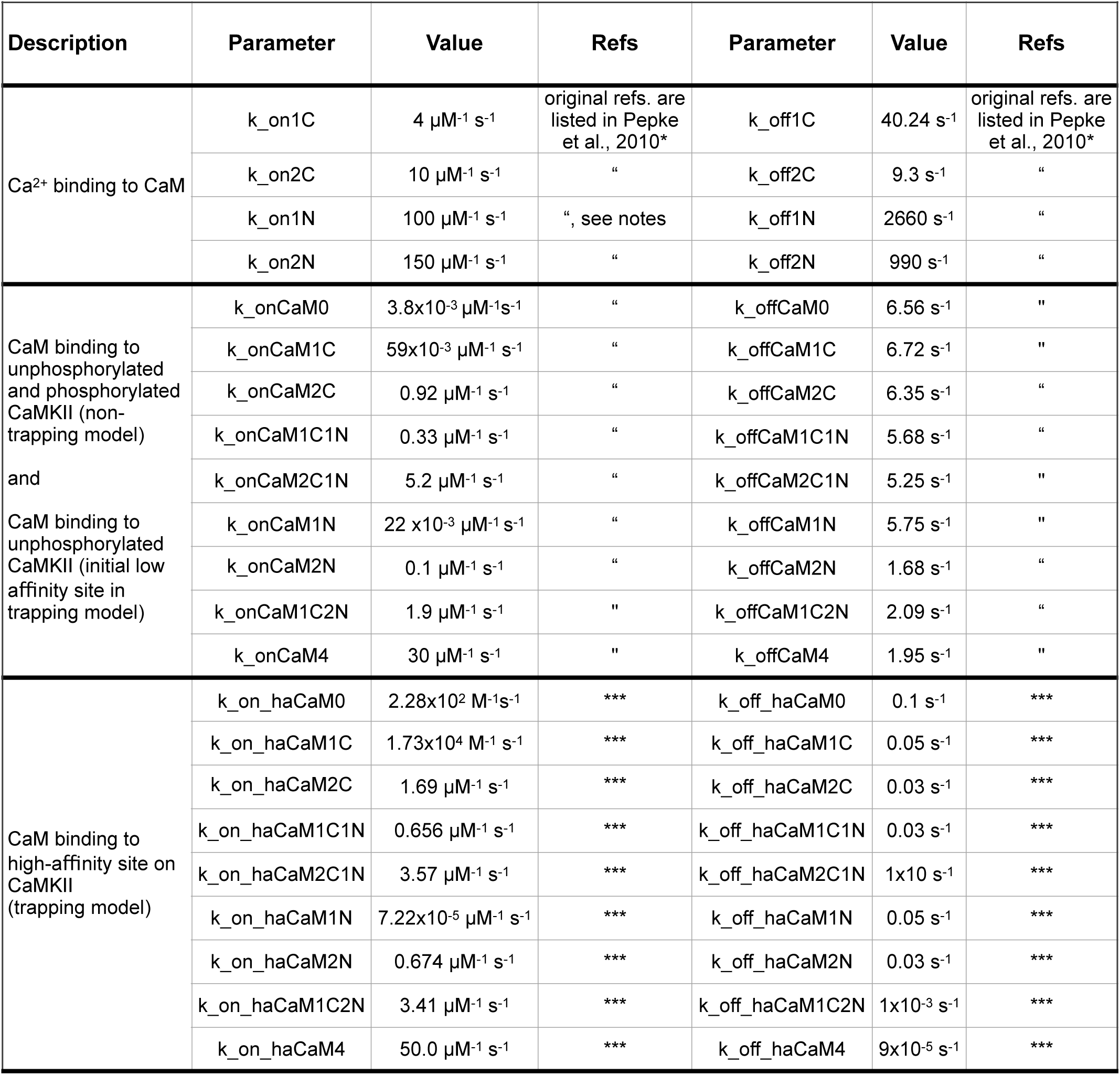
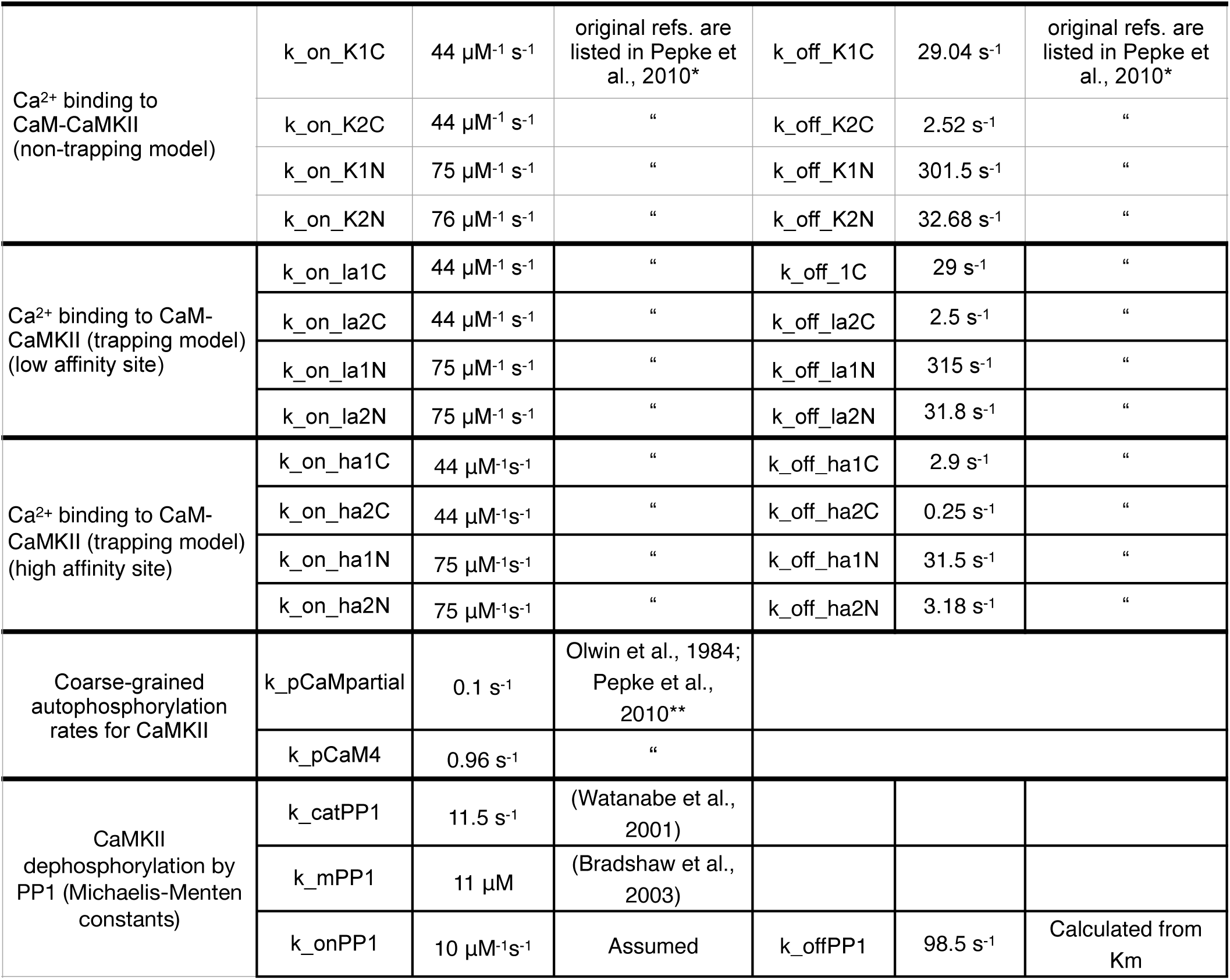
Parameter table with sources of reaction rates for the models of Ca^2+^ and CaM binding to CaMKII. Notes: *The rates from (9) were adjusted to reflect the necessity for detailed balance as described in Pepke et al., 2010. **coarse-grained average phosphorylation rates for CaM-CaMKII with fewer than 4 Ca^2+^ bound and with 4 Ca^2+^ bound were taken from (9) and (10). *** The k_on for CaM4 prior to trapping was taken from (11), and the prolonged k_off for CaM4 in the trapped state was taken from (12). The Remaining k_off rates were estimated from Fig. 3 of (11), and the corresponding k-on rates were calculated from Kd’s calculated to satisfy detailed balance. For the detailed balance calculations in the trapped state, we assumed a 10x higher affinity (10x Kd) of Ca^2+^ for CaM when CaM is bound to the high-affinity site of CaMKII, than when it is bound to the low affinity site. We assumed the higher affinity because the folding of the EF hands around bound Ca^2+^ is stabilized by the binding of CaM to the high-affinity site on CaMKII (i.e. see Fig. 2). We did not use the rapid k_on rate for Ca^2+^ binding to CaM1N and CaM2N calculated in Faas et al. (13) because this rate is not compatible with molecular dynamics measurements (14) or with the rate of removal of water from Ca^2+^ ion (15–17). The k_catPP1 was taken from (5). The k_mPP1 was taken from (18). k_off PP1 was calculated from the equation: *k of f PP*1 = *k on PP*1 * *k m PP*1 − *k cat PP*1. *k_onPP1* was assumed to be 10 µM^-1^ s^-1^.

---

**Figure S1.**
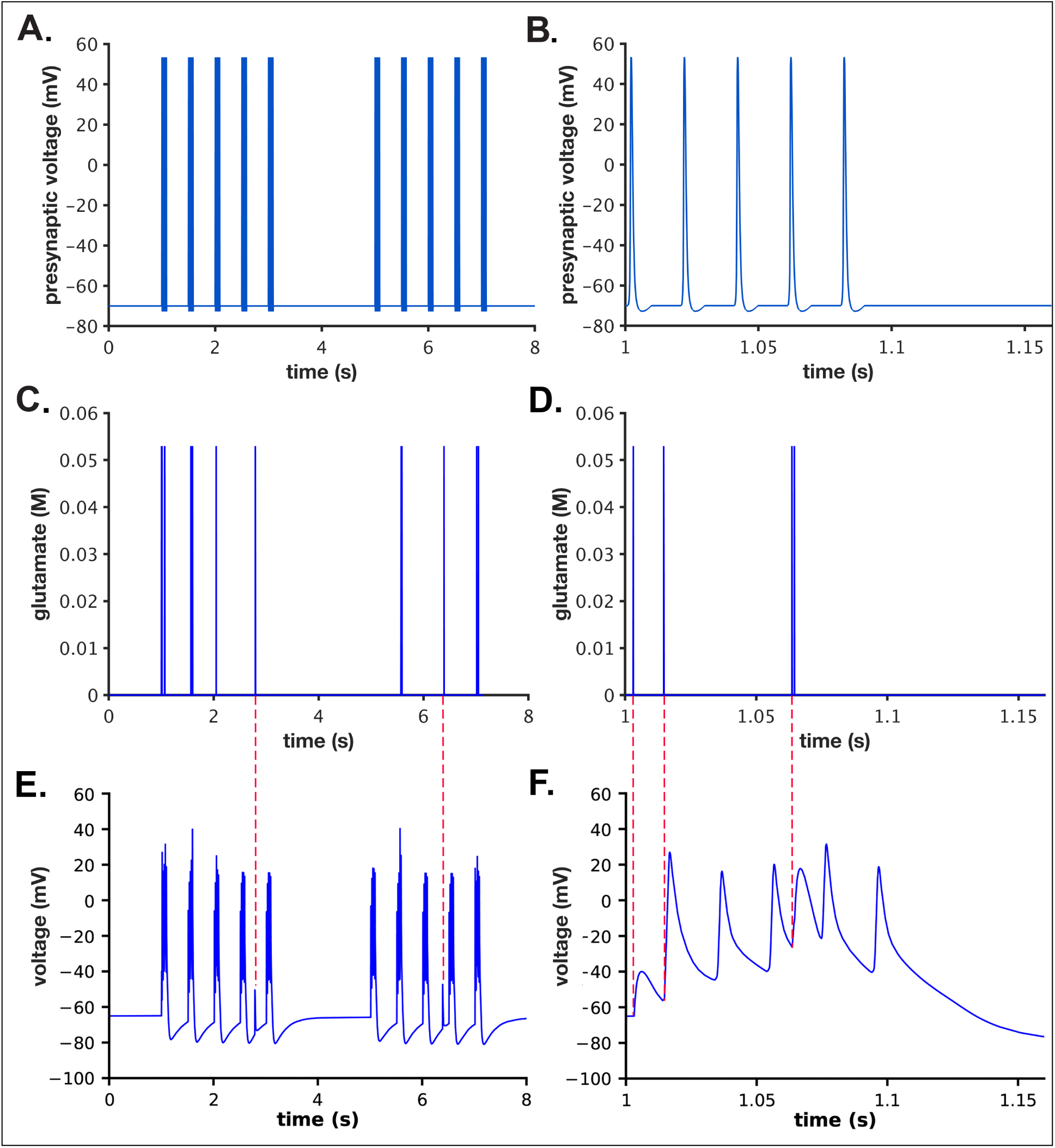
Presynaptic voltage, glutamate release, and postsynaptic voltage during stimulus. A). One example of the fifty different two epoch stimuli of 5 Hz bursts of APs that were applied to the presynaptic bouton as described in the text (Example from a single simulation initiated with seed #50, as described in the text). Each burst was comprised of 5 action potentials delivered at 50 Hz. The stimulus began at 1 sec. The two epochs were separated by 2 secs. B). Data from A. between 1 and 1.15 secs. C). Glutamate concentration in the cleft after release from single vesicles during the stimulus shown in A. D). Data from C between 1 and 1.15 secs. E). Postsynaptic membrane potentials during the single simulation initiated with seed #50. F). Data from E between 1 and 1.15 secs. No glutamate release occurred prior to the third bAP; two release events occurred prior to the fourth bAP; no glutamate release occurred prior to the fifth bAP. Red dashed lines between C. and E. indicate ectopic release events and their associate postsynaptic potentials. Red dashed lines between D. and F. indicate EPSPs generated by glutamate release events depicted in D.

**Figure S2.**
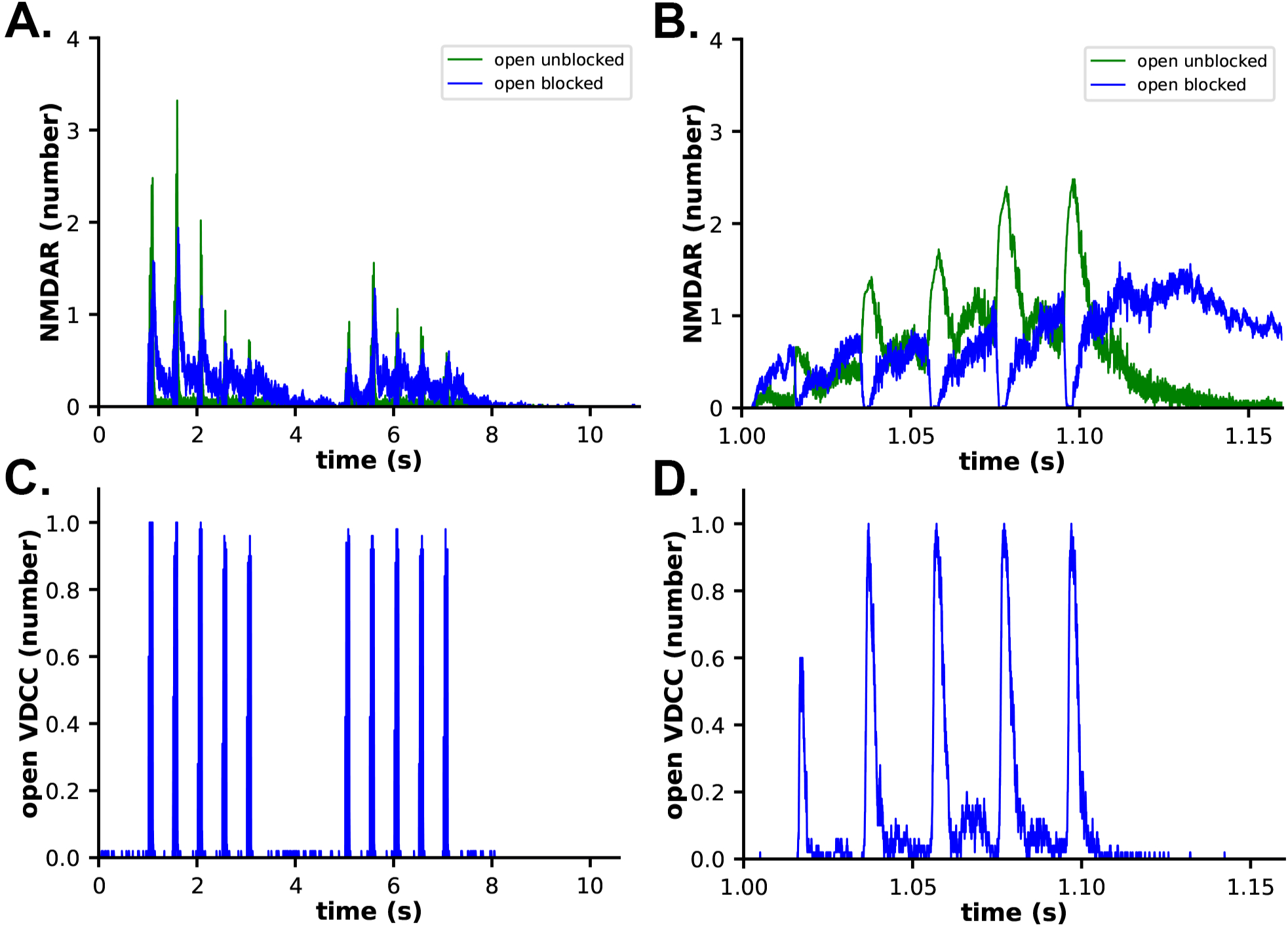
Channel opening kinetics in response to the stimulus shown in. **Figure 5**. A). The number of open, unblocked NMDARs is shown in green, and open, blocked NMDARs in blue. B). Data from A. between 1 and 1.15 s. C). Kinetics of opening of the single VDCC on the spine membrane. D). Data from C. between 1 and 1.15 s. Data are averages of 50 simulations initiated with different random seeds.**Figure S3.** Ca^2+^-bound states of free CaM during the two epoch stimulus.

**Figure S3.**
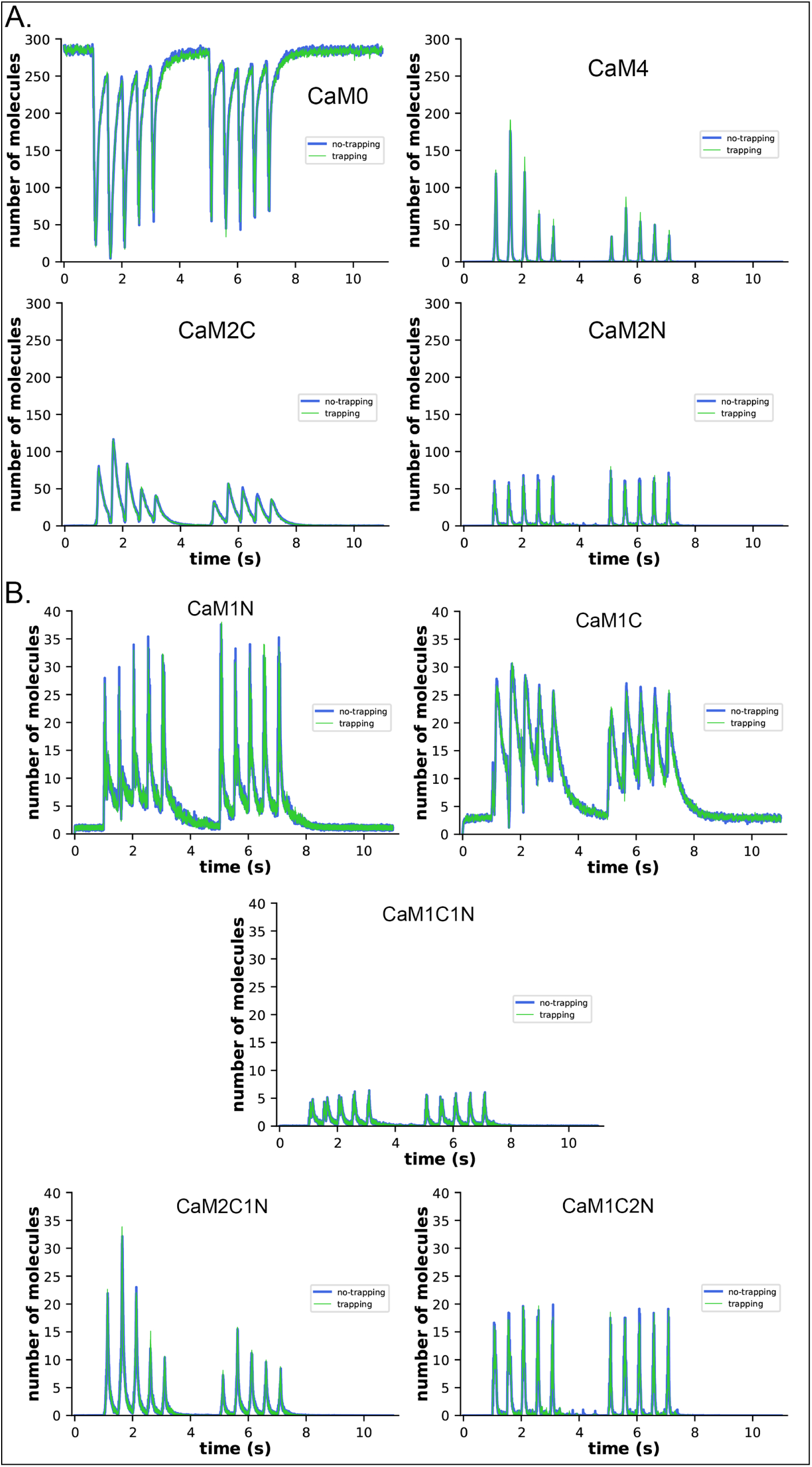
Ca^2+^-bound states of free CaM during the two epoch stimulus. A). Free CaM0, CaM4, CaM2C and CaM2N plotted with ordinate of 300 molecules. B). Free CaM1N, CaM1C, CaM1C1N, CaM2C1N, and CaM1C2N plotted with ordinate of 40 molecules. Blue, no-trapping model; green, trapping model.

**Figure S4.**
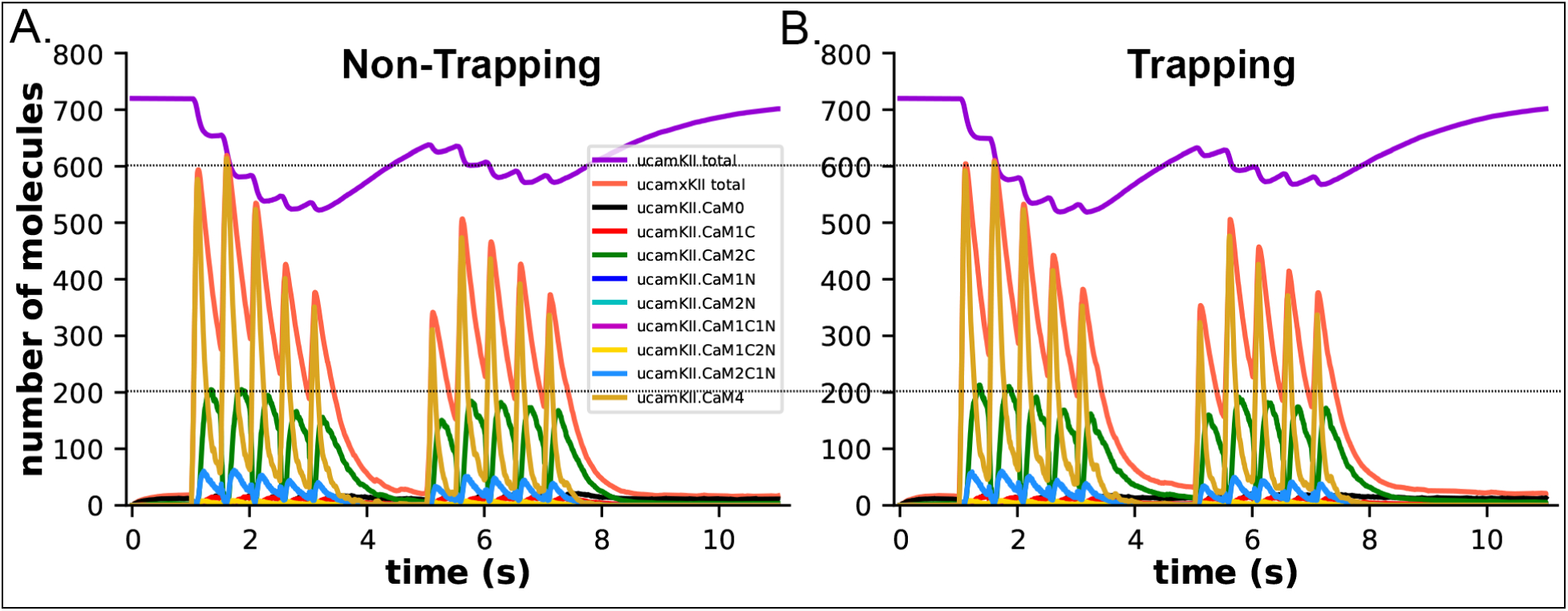
Unphosphorylated CaMKII subunits (ucamkii) and CaM species bound to them during the two epoch stimulus. A). uCaMKII subunits and CaM species bound to them in simulations of the non-trapping model. B). uCaMKII suubnits and CaM species bound to them in simulations of the trapping model. Orange lines indicate the total CaM bound to unphosphorylated CaMKII subunits in each model. There is no significant difference between non-trapping and trapping models in the numbers of CaM species bound to uCaMKII subunits during the stimulus.

**Figure S5.**
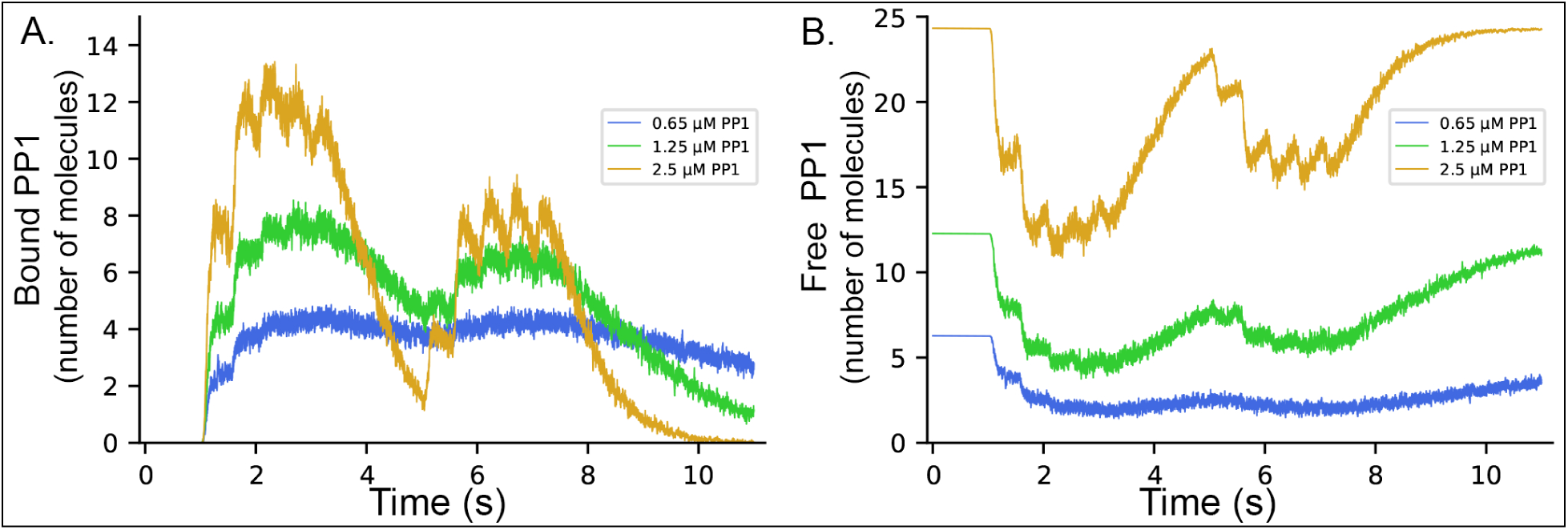
Binding of PP1 to pCaMKII subunits in simulations of the trapping model, in the absence of competition with CaM binding. A). PP1 bound to CaMKII. B). Free PP1. Blue, 0.65 µM PP1; Green, 1.25 µM PP1; Gold, 2.5 µM PP1.

